# Ribosomes in RNA granules are stalled on mRNA sequences that are consensus sites for FMRP association

**DOI:** 10.1101/2021.02.22.432349

**Authors:** Mina N. Anadolu, Jingyu Sun, Senthilkumar Kailasam, Konstanze Simbriger, Teodora Markova, Seyed Mehdi Jafarnejad, Francois Lefebvre, Joaquin Ortega, Christos G. Gkogkas, Wayne S. Sossin

## Abstract

Local translation in neurons is mediated in part by the reactivation of stalled polysomes. Stalled polysomes may be enriched within the pellet of sucrose gradients used to separate polysomes from monosomes. We find that this fraction, isolated from P5 rat brains of both sexes, is enriched in proteins implicated in stalled polysome function, such as the fragile X mental retardation protein (FMRP) and Up-frameshift mutation 1 homolog (UPF1). Cryo-EM analysis of ribosomes in this fraction indicates they are stalled, mainly in the hybrid state. Ribosome profiling of this fraction showed an abundance of footprint reads derived from mRNAs of cytoskeletal proteins implicated in neuronal development and an enrichment of footprint reads on RNA binding proteins. Compared to those usually found in ribosome profiling studies, the footprint reads were more extended on their 3’end and were found in reproducible peaks in the mRNAs. These peaks were enriched in motifs previously associated with mRNAs cross-linked to FMRP in vivo, independently linking the ribosomes in the sedimented pellet to the ribosomes associated with FMRP in the cell. The data supports a model in which specific sequences in mRNAs act to stall translation elongation in neurons.

## Introduction

In neurons, the local translation of mRNAs at distal synaptic sites is essential for neuronal development (1), maintaining the local proteome (2), homeostasis of excitability (3), and synaptic plasticity (4). Local translation requires the transport of mRNAs from the soma to distal sites in a translationally repressed state, followed by their reactivation, either when the mRNA reaches its correct location or after an appropriate stimulus (5). Two major forms of mRNA transport have been defined in neurons: the transport of mRNAs that are repressed at translation initiation and the transport of mRNAs for which elongation has begun but then stalled (6). The mRNAs repressed before elongation lack large ribosomal subunits and are often transported in dedicated mRNA transport particles. In contrast, the mRNAs repressed at elongation are transported as stalled polysomes in RNA granules (6, 7). The local reactivation of translation from mRNAs transported as stalled polysomes can be distinguished from mRNAs blocked at initiation using drugs such as homoharringtonine (HHT) that specifically block the first step of translation elongation, thus blocking translation from mRNA transport particles but not RNA granules (5). Using this tool, several physiological processes have been shown to be supported by initiation-inhibitor resistant protein synthesis, such as the local production of microtubule-associated protein 1B (Map1B) and the induction of a form of long-term depression stimulated by the activation of metabotropic glutamate receptors (mGluR-LTD) in vertebrates, as well as the induction of a type of intermediate term synaptic plasticity in the invertebrate model system *Aplysia californica* (8–10).

Stalled polysomes are transported in large liquid-liquid phase separated structures termed neuronal RNA granules (5). These were first described in oligodendrocytes, where they transported myelin basic protein mRNA to myelin synthesis sites (11). The term neuronal RNA granule was first used to describe a sedimented fraction containing ribosomes and repressed mRNAs lacking initiation factors (7). These large collections of ribosomes can be separated from normal polysomes based on their high density (7,12–15). The proteomic characterization of these structures is consistent with the possibility that they are stalled polysomes (13–15). Indeed, they are enriched in mRNAs such as Map1B that undergo initiation-inhibitor resistant protein synthesis (13).

Stalled polysomes may also be necessary for neuronal development and the regulation of developmentally expressed mRNAs through association with the fragile X mental retardation protein (FMRP), a protein that, when lost, results in the neurodevelopmental disorder Fragile X syndrome (16). UV cross-linking of FMRP to mRNA in neurons showed that FMRP was mainly associated with the coding region of mRNAs (17), consistent with the possibility of FMRP associating with ribosomes. Indeed, FMRP has been shown to be associated with stalled ribosomes in several studies (12,13,17–19). Several mRNA sequences enriched in the regions of mRNAs cross-linked with FMRP have been identified (20, 21) suggesting that specific mRNA sequences may be necessary for determining which mRNAs are recruited to FMRP-containing stalled polysomes. FMRP was also shown to be enriched in RNA granules isolated by sucrose granule sedimentation (13), consistent with the idea that this dense RNA granule fraction contains stalled polysomes.

The mechanism for stalling polysomes during elongation in neurons is unknown. Recently, the nonsense-mediated decay factor UPF1 was implicated in this process. Decreasing levels of UPF1 reduced the levels of initiation-inhibitor resistant protein synthesis, the local production of Map1B and disrupted the induction of mGluR-LTD (8). Since UPF1 is known to be attracted to ribosomes when they reach the stop codon through association with eukaryotic release factors (22), this suggests that stalled mRNAs within RNA granules may be blocked at the release step of translation termination. To test this, we used ribosome profiling, also known as ribosome footprinting, to elucidate the position of ribosomes on mRNAs through sequencing the ribosome protected fragments (footprint reads) after nuclease treatment (23). We took advantage of the presumed enrichment of RNA granules containing stalled polysomes in the pellets of sucrose gradients (13) to create an enriched granule fraction and identified the sites occupied by ribosomes in this fraction. Cryo-EM analysis of the ribosomes in the pellet revealed that most of the ribosomes were stalled in the hybrid position and are loaded with two tRNA molecules, one in the A/P configuration and the second one in a P/E configuration. The footprint reads from these ribosomes were larger than expected and produced reproducible peaks, highly enriched in motifs previously associated with FMRP target mRNAs (20, 21) and consensus sites for m6A modifications in the brain (24). Contrary to our prediction, the footprint reads were not enriched at the stop codon, but instead were slightly biased towards the first half of the open reading frame of transcripts. Cryo-EM analysis of the ribosomes in the pellet revealed that most of the ribosomes were stalled in the hybrid position and are loaded with two tRNA molecules, one in the A/P configuration and the second one in a P/E configuration. We propose a stochastic model in which stalling of ribosomes at specific motifs in mRNAs in neurons attracts FMRP, starting a process of ribosome compaction and RNA granule assembly that allows transport of stalled ribosomes to distal sites in neurons which will later be reactivated for local translation.

## Materials and Methods

### Reagents

Antibodies: rabbit anti-S6 [Cell Signaling #2217], rabbit anti-FMRP [Cell Signaling; #4317], rabbit anti-eEF2 [Cell Signaling; #2332S], rabbit anti-Upf1 [Abcam; ab133564], mouse anti-Stau2 [Medimabs; MM0037-P], rabbit anti-PurA [Abcam; ab79936], mouse anti-hnRNPA2B1 [Novusbio; NB120-6102], mouse anti-SMN [Novusbio; NB100-1936], rabbit anti-IGF2BP1 (ZBP1) [Novusbio; NBP2-38956], rabbit anti-TIA1 [Proteintech; 12133-2-AP], rabbit anti-eIF4E [Cell Signaling; #9742], mouse anti-G3BP [Abnova; H00010146-M01], HRP-conjugated secondary antibodies [ThermoFisher; #31430, #31460].

Enzymes: RNase I [Ambion; AM2294], SuperaseIN [Invitrogen; #AM2696, 20 U/µl], Phosphastase: PNK (T4, NEB; M0201).

Kits: Enhanced chemoluminescence kit [Perkin Elmer; NEL105001EA], Western Blot Stripping Buffer [ZmTech Scientific; S208070], Ribo-Zero Gold (Human/Mouse/Rat) Kit (Illumina; NEBNext^®^ rRNA Depletion Kit (Human/Mouse/Rat)(E6350, NEB); NEXTflex™ Small RNA Sequencing Kit v3 (PerkinElmer, NOVA-5132-06).

Specialized equipment: Bio Rad ChemiDoc digital imager, Biocomp Gradient Master; Agilent Small RNA chip (Agilent Technologies), NovaSeq S1/2 flow cells, Tecnai F20 electron microscope, Gatan Ultrascan 4000 4 k × 4 k CCD Camera System. Density Gradient Fractionation System [Teledyne ISCO; BR-188; Brandel] with Syringe Pump [Syr-101; Brandel], Fraction Collector [Foxy R1; Teledyne ISCO] with UA-6 Absorbance Detector [Brandel].

**Biological Resources:** Sprague Dawley Rats (Charles River Laboratories)

**Computational Resources**: See **Riboseq and RNAseq data analysis**

**Statistical analysis**: ANOVAs were used to determine significant enrichment and abundances of particular sets of mRNAs in the stalled polysomes compared to the overall mRNAs. Significance of Motifs and GO terms were determined by the programs utilized for this task (Homer, gprfilerR), limma (see Riboseq and RNAseq data analysis section).

### Purification of RNA Granules

RNA Granules were purified, from whole brain homogenate harvested from five-day-old (P5) Sprague Dawley rats of both sexes, using a protocol adapted from a previous study (13). Five P5 rat brains were homogenized in RNA Granule Buffer (20 mM TRIS-HCl pH 7.4 [Fisher; BP152-1], 150 mM NaCl [Fisher; BP358-212], 2.5 mM MgCl_2_ [Fisher; M33-500]) supplemented with 1 mM DTT [Sigma; D9163], 1 mM EGTA [Sigma; E8145], EDTA-free protease inhibitor [Roche; 04693132001]. Note that cycloheximide [Abcam; ab120093] was not added to the homogenization buffer unless explicitly stated. Homogenate was centrifuged 15 minutes in a Thermo Scientific T865 fixed angle rotor at 6117 x g at 4°C to spin down cellular debris. The supernatant was treated with 1% IGEPAL CA-630 [Sigma; I8896] for 5 minutes at 4°C on a rocker. The sample was then loaded onto a 2 ml 60% sucrose [Calbiochem; 8550] cushion (dissolved in supplemented RNA Granule Buffer) in a Sorvall 36 ml tube [Kendro; 3141, Thermo Scientific], filled to top with additional RNA Granule Buffer and centrifuged for 2 hours in a Thermo Scientific AH-629 swing-bucket rotor at 56660 x g at 4°C to achieve the polysome pellet. The pellet was re-suspended in RNA Granule Buffer, gently dounced and loaded over a 15-60% linear sucrose gradient (gradient was made with RNA Granule Buffer) that was prepared in advance using a gradient maker [Biocomp Gradient Master] and centrifuged for 45 minutes at 56660 x g at 4°C in an AH-629 swing bucket rotor. Fractions of 3.5 ml were then collected from the top, and the remaining pellet was rinsed once and then resuspended using RNA Granule Buffer. For experiments measuring UV absorbance, sucrose gradients were centrifuged using a SW40 rotor (Beckman Coulter) and fractionated using an ISCO density gradient fractionation system and optical density was continuously recorded at 254  nm and fractions were collected with a FOXO JR Fractionator (Teledyne ISCO). For some experiments, such as for Electron Microscopy, the resuspended pellet was used directly, or treated with salt and nuclease to break up the ribosome clusters into monosomes (see below). For some experiments, such as ribosome footprinting (see Nuclease and Salt Treatments below), the fractions were precipitated overnight at −20 °C by adding 7 ml of chilled 100% ethanol. The precipitated samples were then centrifuged for 45 minutes at 2177 x g at 4°C in an Eppendorf 5810/5810 swing bucket rotor before collection using RNA Granule Buffer.

### Nuclease and Salt Treatments

We treated the pellet fraction with salt and nuclease to break up the stalled polysomes compacted in the RNA Granule into monosomes for ribosome footprint analysis. The pellet was incubated with RNA granule buffer containing 400 mM NaCl for 10 minutes at 4°C on a rocker(13). Before nuclease treatment, the NaCl concentration was reduced back to 150 mM by diluting the sample with a NaCl-free RNA Granule Buffer. The sample was then treated with 100 U of RNase I [100 U/µl; Ambion AM2294, Thermo Fisher] for 30 minutes at 4°C on a rocker. The nuclease was quenched with 100 U of SuperaseIN [20 U/µl; Invitrogen #AM2696, Thermo Fisher], and the samples were re-run on a fresh 15-60% sucrose gradient to separate mononsomes. Fraction 2 was precipitated overnight at −20 °C by adding 7 ml of chilled 100% ethanol. The precipitated samples were then centrifuged for 45 minutes at 2177 x g at 4°C in an Eppendorf 5810/5810 swing bucket rotor before collection using RNA Granule Buffer.

### Transmission and Cryo-Electron Microscopy

In experiments where samples were imaged by negative staining, the untreated pellet fraction, and the fractions after treatment with either high-salt or nuclease or both, were deposited on the EM grids. The ribosome concentration of each sample was adjusted to ∼80 ng/μl (∼25 nM) using RNA Granule Buffer before applying them to the grids. In the case of the sample treated with both nuclease and high salt the concentration of sample applied to the grid was 9.2ng/μl (2.9 nM). We used 400-mesh copper grids freshly coated with a continuous layer of thin carbon for these experiments. Grids were glow-discharged at 15  mA for 15  seconds and then floated on a 5-µl drop of the diluted sample for 2  min. Excess of sample was blotted away with filter paper (Whatman #1), and to stain them, they were subsequently floated in a 5-µl drop of a 1% uranyl acetate solution for 1  min. Excess of stain was blotted away, and the grids were dried on air and stored in regular grid boxes. The EM images were acquired on a Tecnai F20 electron microscope operated at 200 kV using a room temperature side entry holder. Images were collected in a Gatan Ultrascan 4000 4 k × 4 k CCD Camera System Model 895 at a nominal magnification of 60,000x. Images produced by this camera had a calibrated pixel size of 1.8 Å/pixel. The total electron dose per image was ∼50 e-/Å2. Images were collected using a defocus of approximately −2.7 µm. Images were prepared for figures using the Adobe Photoshop program.

For samples imaged by cryo-electron microscopy (cryo-EM), the pellet fraction was treated with nuclease before being deposited on the cryo-EM grids. The ribosome concentration in the sample applied to the grid was 160 nM. Cryo-EM grids (c-flat CF-2/2-2C-T) used for these samples were washed in chloroform for two hours and treated with glow discharged in air at 15 mA for 20 seconds. A volume of 3.6 μL was applied to the grid before vitrification in liquid ethane using a Vitrobot Mark IV (Thermo Fisher Scientific Inc.). The Vitrobot parameters used for vitrification were blotting time 3 seconds and a blot force +1. The Vitrobot chamber was set to 25 °C and 100% relative humidity.

Cryo-EM datasets were collected at FEMR-McGill using a Titan Krios microscope at 300 kV equipped with a Gatan BioQuantum K3 direct electron detector. The software used for data collection was SerialEM (25). Images were collected in counting mode according to the parameters described in Supplementary Table 1 (see below).

To calculate the cryo-EM structures, cryo-EM movies obtained in the Titan Krios were corrected for beam-induced motion using RELION’s implementation of the MotionCor 2 algorithm(26). CTF parameter estimation was done using the CTFFIND-4.1 program (27). The remaining processing steps were done using RELION-3 (28). Individual particles in the images were identified using auto-picking. These particles were extracted and subjected to 2 cycles of reference-free 2D classification to remove false positives from the auto picking process. The cleaned dataset was subjected to 3 layers of 3D classification. In each layer, each class was split into three new classes. The initial 3D reference used for the 3D classification was a 60 Å low pass filtered map of the 80S human ribosome created from EMD-2938 (29). The 3D classifications did not used masks. To speed up the computer calculations, the 2D and 3D classifications were performed using particle images binned by 4. However, we use full-size images (1.09 Å/pixel) in the refinement steps. Particles assigned to maps representing the same conformation were pooled together and used for 3D auto-refine. The numbers of particles assigned to each class and included in each refinement are described in Supplementary Table S1. Refinement was performed in four stages: In the first stage, the 3D auto-refine was performed without a mask. The resulting map was used to create a mask and was also used as the initial reference for a second stage refinement that included one additional 3D auto-refine cycle. Masks were created with ‘relion_mask_create’ command extending the binary mask by four pixels and creating a soft edge with a width of four pixels. The initial threshold for binarization of the mask varied depending on the structure. In the third step, we used the output of the last 3D auto-refine job as the input for CTF refinement. In this process, we selected ‘Estimate (anisotropic) magnification’ as yes in a first cycle. In the second cycle of CTF refinement, we selected ‘Perform CTF parameter fitting’ as Yes and selected ‘Fit defocus’ as ‘Per-particle’ and ‘Fit astigmatism as ‘Per-micrograph.’ We selected ‘No’ for ‘Fit B-factor’, ‘Fit phase-shift’, ‘Estimate trefoil’ and ‘Estimate 4th order aberrations. However, we selected ‘Yes’ for ‘Estimate beam tilt, as our data was collected using this approach. We also selected ‘Yes’ for ‘Estimate trefoil’ and ‘Estimate 4th order aberrations’. In the fourth step, we used the output from the CTF refinement job to perform Bayesian polishing to correct for per-particle beam-induced motion before subjecting these particles to the last cycle of 3D auto-refine. Bayesian polishing was performed using sigma values of 0.2 Å/dose, 5,000 Å and 2 Å/dose for velocity, divergence and acceleration, respectively. The map from the last cycle of the 3D auto-refine was used to perform multi-body refinement by dividing the 80S structure into 2 major bodies (body1: 60S, body2: 40S). Sharpening of the final cryo-EM maps was done with RELION (28). The average resolution for each cryo-EM map obtained from the multi-body refinement was estimated by gold-standard Fourier shell correlation. Resolution estimation is reported using a FSC threshold value of 0.143. Local resolution analysis was done with RELION (28). In each class, the cryo-EM maps of the 40S and 60S subunits derived from the multi-body refinement were merged into a cryo-EM map for the 80S ribosome using “vop add” command in Chimera. Cryo-EM map visualization was done by UCSF Chimera (30) and Chimera X (31, 32).

### Immunoblotting and Quantification of Enrichment

For immunoblotting, SDS sample buffer was added to each sample before loading onto a 10% or 12% acrylamide gel. The resolved proteins were either stained with Coomassie brilliant blue [Fisher; 821616] or transferred onto a 0.45 µm nitrocellulose membrane [Bio Rad; 1620115] for immunoblotting. The membranes were blocked with 5% BSA [Sigma; 9647] in Tris Buffered Saline with Tween (TBS-T [Tris: Fisher, BP152-1; NaCl: Fisher, BP358-212; Tween: Fisher, BP337]) before incubation with primary antibodies (see reagents) all at 1:1000 dilution.

Membranes were washed with TBS-T after incubation. Detection was done using HRP-conjugated secondary antibodies [ThermoFisher; #31430, #31460] before ECL [Perkin Elmer; NEL105001EA] reaction and imaging using the Bio Rad ChemiDoc digital imager. For quantification of RBP enrichment, membranes were stripped with a Western Blot Stripping Buffer [ZmTech Scientific; S208070] and re-probed with rabbit anti-S6 [Cell Signaling #2217] antibody, followed by detection with HRP-conjugated secondary antibodies [ThermoFisher; #31430, #31460].

Quantification of signal intensity was done using ImageJ software. We selected full lane ROIs and quantified single bands corresponding to the observed kDa size of each protein. We then used the corresponding anti-S6 signal intensity to quantify the amount of examined protein per S6 ribosomal protein for each fraction. For each experiment the protein/S6 value was normalized to the protein/S6 value from the starting material. Following this normalization, biological replicates were averaged. For salt and nuclease experiments, we calculated the proportion of Granule Fraction Polysomes (Pellet) digested into monosomes (Fraction 2) by doing the following calculation: Fraction 2/(Fraction 2 + Pellet).

### Footprint sequencing library construction and sequencing

Fraction 2 from the monosome purification was centrifuged at 4,444 x g at 4 °C for 60 min, the isopropanol taken off, the pellet air-dried, and re-suspended in 10 mM Tris-HCl pH 7. We then proceeded with a standard hot phenol-acid extraction (Acid Phenol:Chloroform mix 125:24:1, Invitrogen, AM9722). The samples were resuspended in nuclease-free water and quantified on a Spectrophotometer. All samples were depleted of ribosomal RNA using the Ribo-Zero Gold (Human/Mouse/Rat) Kit (Illumina) (replicates 1-3 and cycloheximide treated) or the NEBNext^®^ rRNA Depletion Kit (Human/Mouse/Rat)(E6350, NEB) (replicates 4-5 and no salt treatment) The footprint samples were size-selected using the 17 and 34 nt markers as a guide on a 15% TBE-Urea polyacrylamide gel (Thermo Scientific), while total RNA samples were randomly heat fragmented by incubating at 95 °C in an alkaline fragmentation solution (50 mM NaCO_3_ pH 9.2, 2 mM EDTA) for 40 min, yielding comparably sized fragments to footprints. All samples were de-phosphorylated using PNK (T4, NEB). The quality and concentration of samples was assessed by running an Agilent Small RNA chip (Agilent Technologies), and sequencing libraries were generated using the NEXTflex™ Small RNA Sequencing Kit v3 (PerkinElmer, NOVA-5132-06), according to the manufacturer’s instructions. Samples were balanced and pooled for sequencing with Edinburgh Genomics on NovaSeq S1/2 flow cells, yielding 50 bp paired-end reads.

### Riboseq and RNAseq data analysis

Adaptor sequences and low-quality score containing bases (*Phred score < 30*) were trimmed from reads using Cutadapt v2.8 (-*j 8 -u 4 -u −4 -Z -o*) (33). Noncoding RNAs were removed by custom scripts following mapping these contaminant reads using Bowtie2 v2.3.5 (*--phred33 -- very-sensitive*) (34). The unmapped reads were aligned to reference rat genome (Rnor_6.0.94) using *STAR v2.7.3a* (*35*)with options previously described (36) (--*twopassMode Basic -- twopass1readsN-1 -- seedSearchStartLmax 15 --outSJfilterOverhangMin 15 8 8 8 -- outFilterMismatchNoverReadLmax 0.1*). QuantMode with STAR was used to obtain genomic and transcript coordinates (35). Assigning RPF reads to genomic features (CDS, UTRs) was based on Genome annotation (Rnor_6.0.94). Only a single transcript isoform, with the highest APPRIS score (37), was considered per gene. All raw reads from these experiments have been deposited at the GEO dataset at NCBI (number to be confirmed).

Raw counts obtained using *featurecounts v2.2.0* (38) were analyzed using the R package *limma* (39). From this package, transcript abundance was obtained for riboseq and RNAseq data in reads per kb of mRNA (RPKM). Enrichment was determined for each transcript by dividing riboseq RPKM by RNAseq RPKM. False discovery rates (FDR) and p values were determined for enrichment in the package, and nominal p-values were corrected for multiple testing using the Benjamini-Hochberg method.

Gene Ontology (GO) enrichment analysis was performed with *gProfileR* (40). Enrichment p-values were based on a hypergeometric test using the set of known Rat genes as background. Sample correlation based on normalised read count was obtained using the R package *limma* (39). We used Ribowaltz (41) to assign length dependant P-site correction and periodicity assignment to each read.

### Identification of Consensus Peaks

Sites enriched with ribosomal footprints were identified using normalized riboseq profiles using peak identification function within *IDPmisc,* a R package (42). Peak width of minimum 18 nt and peak height above the mean peak height within the transcript was used as the criteria to define a region with significant enrichment of RPFs. Only peaks region that overlap (at least 90% nt overlap) in at least 3 out of 5 samples were considered using BEDtools v2.29.2 (43)

### Motif analysis

The peak regions were scanned for known human RNA-binding protein motifs using the *FIMO* program (44) which is part of MEME Suite (45). Only search results with a p-value less than the threshold of 0.5 were considered. De novo motif finding was performed using peak from RPFs. *HOMER* tool (46) was used for this analysis (*findMotifs.pl-rna)*. Background sequences were randomly selected from transcripts with no peaks.

## Results

### Isolation of the granule fraction

We used a previously described protocol (13) for sucrose gradient sedimentation that involves a high-velocity spin over a 60% sucrose pad to enrich for all polysomes, followed by short spin on a 15-60% sucrose gradient to separate polysomes from a granule fraction that may represent stalled ribosomes that are transported in neuronal RNA granules (Fig. 1A). As previously described (13), the short sedimentation allows for clear separation of two populations of ribosomal proteins as can be seen from Coomassie staining and immunoblotting for the ribosomal protein S6 (Fig. 1B). The ribosomal proteins peak in fraction 5-6 and very few ribosomal proteins are found in fractions 8-9 (Fig. 1B) in contrast with the large number of ribosomal proteins in the pellet fraction (Fig. 1B). A UV absorption plot also confirms that very few A254 absorbing structures are present before the pellet (Fig. 1C). EM of the pellet shows ribosomal clusters in transmission EM pictures (Fig. 1D). Counting of ribosomes in the clusters from the pellet or granule fraction (GF) and the peak of the polysome fraction (PF)(Fractions 5-6) using transmission EM micrographs revealed no difference in the number of ribosomes/cluster between the two fractions suggesting that the granule fraction is not simply made up of larger polysomes (10 ± 4.2 ribosomes in the GF (n=219 clusters, 14 micrographs, 2 preparations) and 10 ± 5.2 ribosomes in the PF (N=240 clusters, 14 micrographs, 2 preparations).

**Figure 1.**
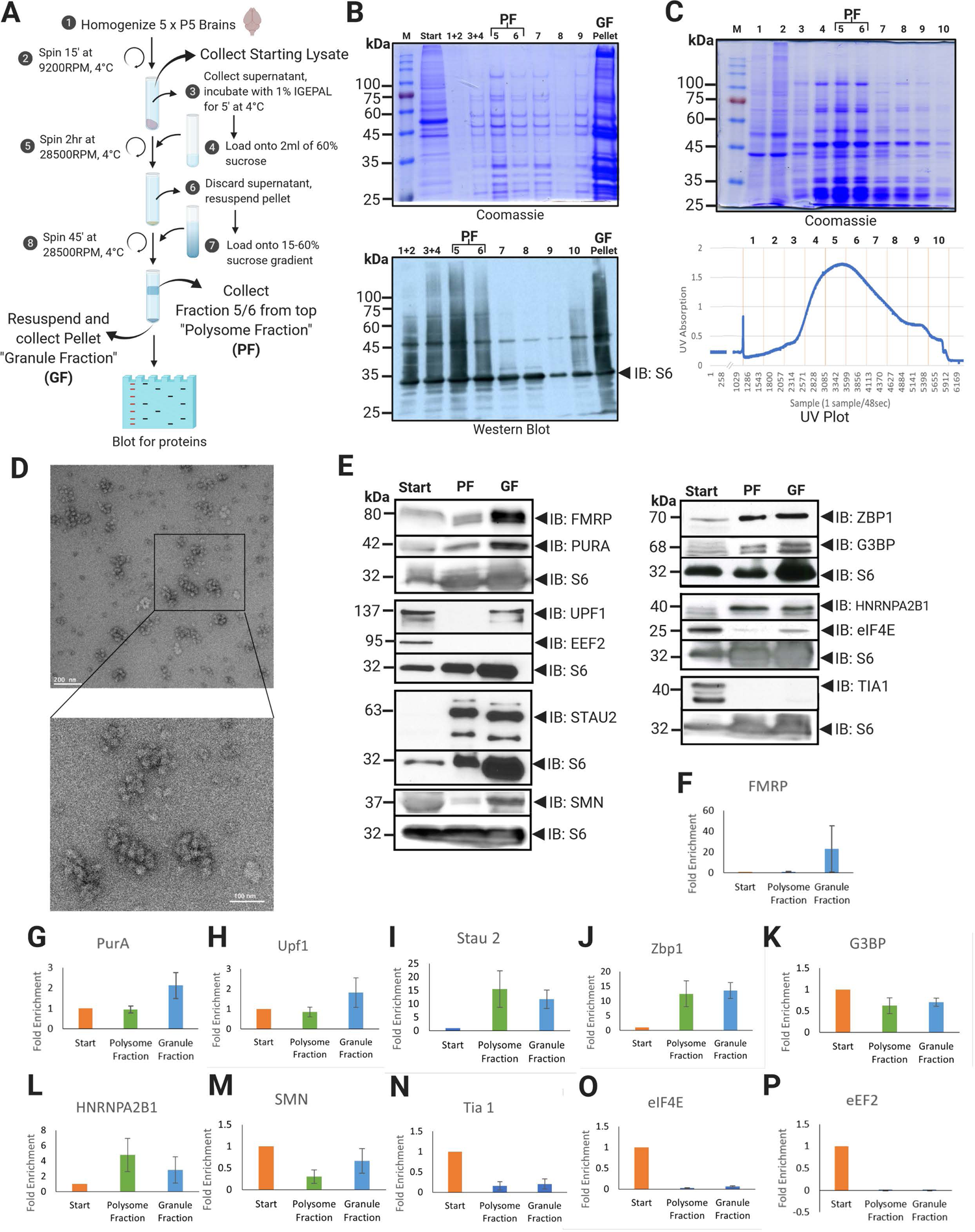
Isolation of a pellet enriched in compacted ribosomes. (A) Summary of protocol for isolating the pellet/granule fraction (GF) from P5 rat whole brain homogenate using sucrose gradient fractionation. (B) Top: SDS-page stained with Coomassie brilliant blue showing enrichment of the characteristic distribution of ribosomal proteins in Fraction 5-6 (Polysome Fraction; PF) and Pellet (GF); Bottom: Immunoblot analysis (from a separate purification) of S6 ribosomal protein showing peaks in Fractions 5-6 (PF) and Pellet (GF); Lanes are described on top (M for molecular weight marker). (C) Top: SDS-page stained with Coomassie brilliant blue showing the distribution of proteins from Fraction 1 to Fraction 10, excluding the pellet. Bottom: UV absorption plot (A254) of the same experiment collected fraction 1-10 showing enrichment of polysomes in the PF. (D) Negative stained electron micrograph of the GF shows clusters of ribosomes; inset showing close up of clustered ribosomes (E) Immunoblot analysis of starting material, Polysome Fraction (PF; Fraction 5/6) and Granule Fraction (GF; Pellet) stained for RBPs implicated in RNA granules and stalled polysomes and other factors: FMRP, PURA, Upf1, Stau2 and eEF2, ZBP1, G3BP, hnRNPA2B1, eIF4E, SMN, and Tia-1. One representative blot for each RBP is shown and each blot is normalized to the S6 staining from that blot and the S6 blot is shown underneath each separate blot. (F)-P) Quantification of Western Blots. The fold enrichment compared to starting material normalized to levels of S6 (see methods) is shown for each RBP: FMRP (N=3), PURA (N=3), Upf1 (N=6), Stau2 (N=4), eEF2 (N=3), ZBP1 (N=4), G3BP (N=3), hnRNPA2B1 (N=3), eIF4E (N=4), SMN (N=3), and Tia-1 (N=4). Error bars represent SEM.

One possibility is that the GF represents aggregated polysomes as opposed to a denser fraction of ribosomes. If the GF represented aggregates of regular polysomes, one would expect to observe equal distributions of RNA binding proteins (RBPs) in both fractions. To examine this, we performed western blot analysis on the starting material, PF and GF, examining RNA binding proteins (RBPs) known to be differentially distributed in RNA granules/stalled polysomes. For quantification, we standardized the levels of the proteins to the levels of the ribosomal protein S6 to normalize for the number of ribosomes in each fraction. FMRP has been linked to stalled polysomes in several studies (9,13,17,18,47) and was highly enriched in the GF (Fig. 1E, F). Similarly, Pur-alpha (PURA), a protein that marks RNA granules in neurons (13, 15) and whose loss leads to neurodevelopmental disorders (48), was enriched in the GF compared to the PF (Fig. 1E, G). UPF1, the key component of the nonsense-mediated decay pathway (22), has been shown to play an independent role in the formation of stalled polysomes in neurons (8) is also enriched in the GF compared to the PF (Fig. 1E, H). UPF1 interacts with Staufen 2 (STAU2), and this interaction is important for the formation of stalled polysomes (8). However, while STAU2 was present in the GF, it was not enriched compared to the PF (Fig. 1E, I). All of the other RBPs (ZBP1, G3BP1, hnRNPA2B1, SMN) that have also been associated with RNA granules in previous studies (14, 49) showed similar results to STAU 2 (Fig 1E, J-M), suggesting that many RBPs are equally distributed between the two fractions and emphasizing the enrichment of FMPR, PURA and UPF1 in the GF, specifically. In contrast, TIA, an RBP particularly implicated in stress granules (50), but not present in RNA granule proteomics (13–15) was not enriched in either of the two fractions (Fig. 1E, N). Neither the polysome nor the pellet fraction was enriched for eIF4E, consistent with the lack of ribosomes in the process of initiation (Fig. 1E, O). Interestingly, we were unable to detect the presence of the eukaryotic elongation factor 2 (eEF2) in the polysome or the granule fraction (Fig. 1E, P). Given the proposed role for eEF2 phosphorylation in the reactivation of stalled polysomes (4, 10), the absence of eEF2 suggests that this may be due to a requirement for eEF2 release from normal translating polysome to restart stalled polysomes.

Overall, the protein composition of the GF (pellet) demonstrates distinctions in the molecular components of ribosomes compared to the PF (fractions 5-6), suggesting a distinct subset of ribosomes are found in the pellet, either due to difference in their density, or an increased propensity for aggregation.

### Cleavage of compacted ribosomes into monosomes

Ribosome profiling is based on nuclease digestion of mRNA that is not protected by ribosomes, isolation of monosomes containing the protected mRNA, followed by library construction and sequencing. Compacted ribosomes are resistant to cleavage by nucleases (17). Based on the previous observation that high-salt conditions cause the unpacking of the compacted ribosomes (13), we treated the granule fraction with 400 mM of sodium chloride for 10 minutes before dilution to reduce the salt concentration back to physiological levels (150 mM) followed by incubation with RNase I (Fig. 2A). We found that this treatment could cleave the compacted ribosome clusters to monosomes (Fig. 2B-F). The effect of the high-salt and nuclease treatments were visualized using negative staining electron microscopy (EM) (Fig 2B). Samples either untreated or treated with the high-salt buffer, nuclease (low concentration; 0.5 µl: 50 U) or both were deposited on the EM grids and negatively stained. We observed that the high-salt and nuclease treatments, when applied separately, induced partial unpacking of the ribosome clusters. Only when both treatments were applied to the sample did the EM images show that the ribosome clusters had dissociated into monosomes (Fig 2B).

**Figure 2.**
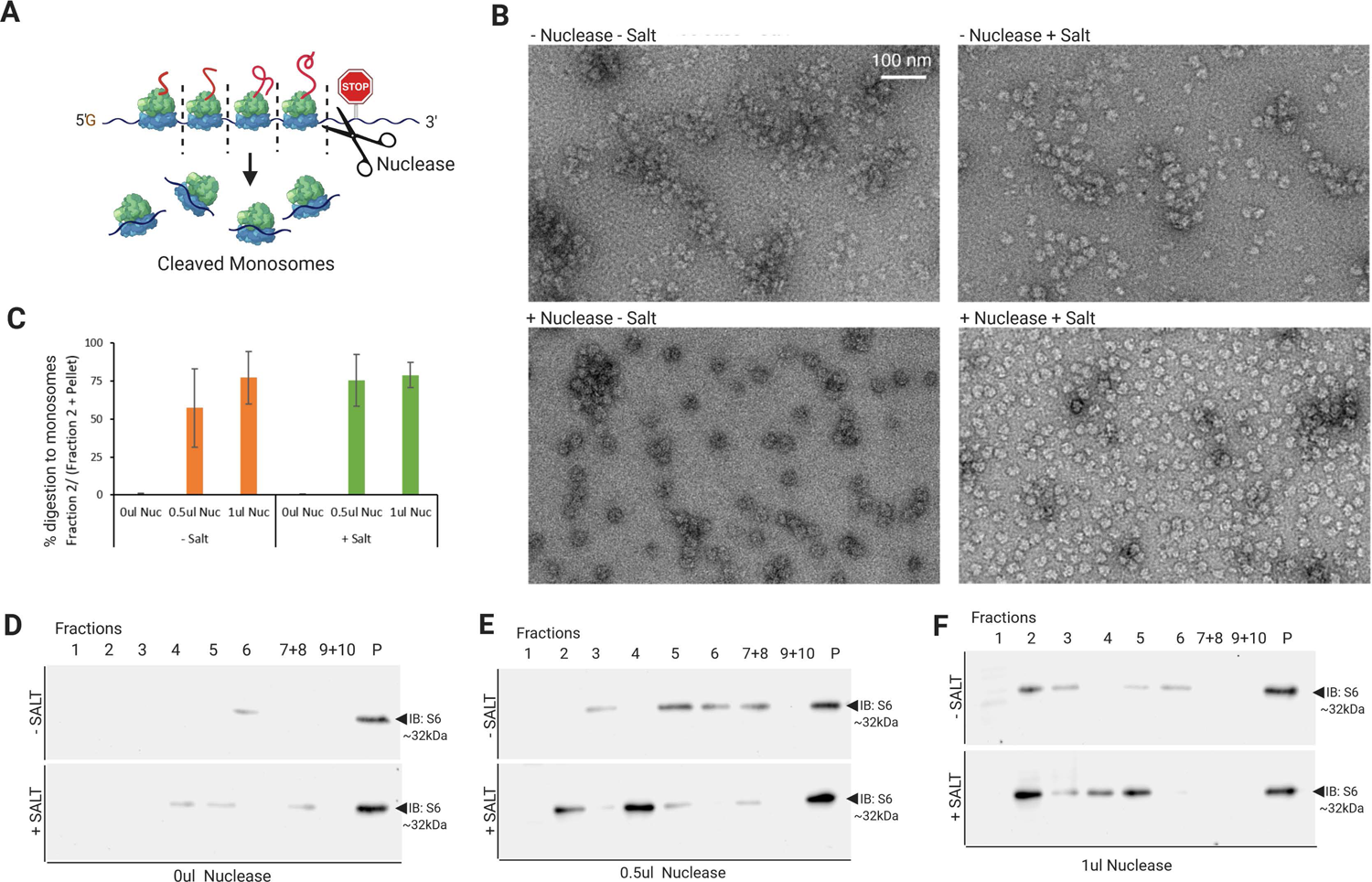
Cleavage of compacted stalled polysomes into monosomes. (A) Schematic of nucleases cleaving polysomes into monosomes. (B) Electron micrographs of negatively stained pellet fraction following treatment with RNase I, with and without pre-treatment with Salt (-Nuclease -Salt: TOP LEFT; -Nuclease +Salt: TOP RIGHT; +Nuclease – Salt: BOTTOM LEFT; +Nuclease +Salt: BOTTOM LEFT). Scale bar represents 100 nm. Note that EM images represent the pellet fraction treated with Nuclease and Salt before the second sucrose gradient and not the purified monosomes in Fraction 2 (C) Quantification of digestion represented as the ratio of Fraction 2/(Fraction 2 + Pellet), N=4 biological replicates, error bars represent SEM (D-F) Western Blot analysis for S6 ribosomal protein in sucrose gradient fractions. The GF was resuspended, treated with 0ul (D), 0.5ul (E) and 1ul (F) RNase I, with or without pre-treatment of 400 mM NaCl (-SALT: TOP; +SALT: BOTTOM), followed by a 15-60% sucrose gradient run for 45 min (see Methods)

We also examined cleavage by re-sedimentation after cleavage. After RNase digestion, we re-sedimented the fractions on a sucrose gradient and measured the movement of S6 ribosomal protein immunoreactivity from the GF to fraction 2 of the sucrose gradient (Fig. 2C-F). We observed that the high-salt treatment improved the digestion by nuclease at low nuclease concentrations. However, some preparations of nuclease led to complete cleavage even without the need for salt (Supp Fig. 1). At the higher concentration of nuclease (1 µl: 100 U), there was no difference in digestion in the presence or absence of salt (Fig 2C; Supp Fig. 1), consistent with previous results suggesting that high concentrations of nuclease can cleave compacted ribosomes (17).

### Cryo-EM analysis of ribosomes reveals ribosomes are stalled in the hybrid position

To establish whether the ribosomes in the RNA granules are paused at the start of translational elongation or after elongation has started and then stalled, RNAase I-treated RNA granules were deposited in cryo-EM grids and vitrified for analysis using single-particle approaches. The 400 mM sodium chloride treatment of the RNA granules was eliminated for these samples to prevent dissociation of stalling factors potentially bound to the stalled ribosomes. Individual ribosomes in the cryo-EM micrographs were selected and subjected to 2D and 3D classification. We found that two classes of the 80S ribosomes co-existed in the RNA granules. Class 1 was more populated and contained 85% of the particles in the dataset. The cryo-EM map for class 1 refined to 3Å resolution (Suppl. Fig. 2A) and exhibited tRNA molecules in hybrid A/P and P/E state (Fig. 3A). Particles in class 2 represented the 15% remaining of the population, and their cryo-EM map refined to 2.6Å resolution (Suppl. Fig2B), revealing a tRNA in the P-site (Fig. 2B), but an empty A site.

**Figure 3.**
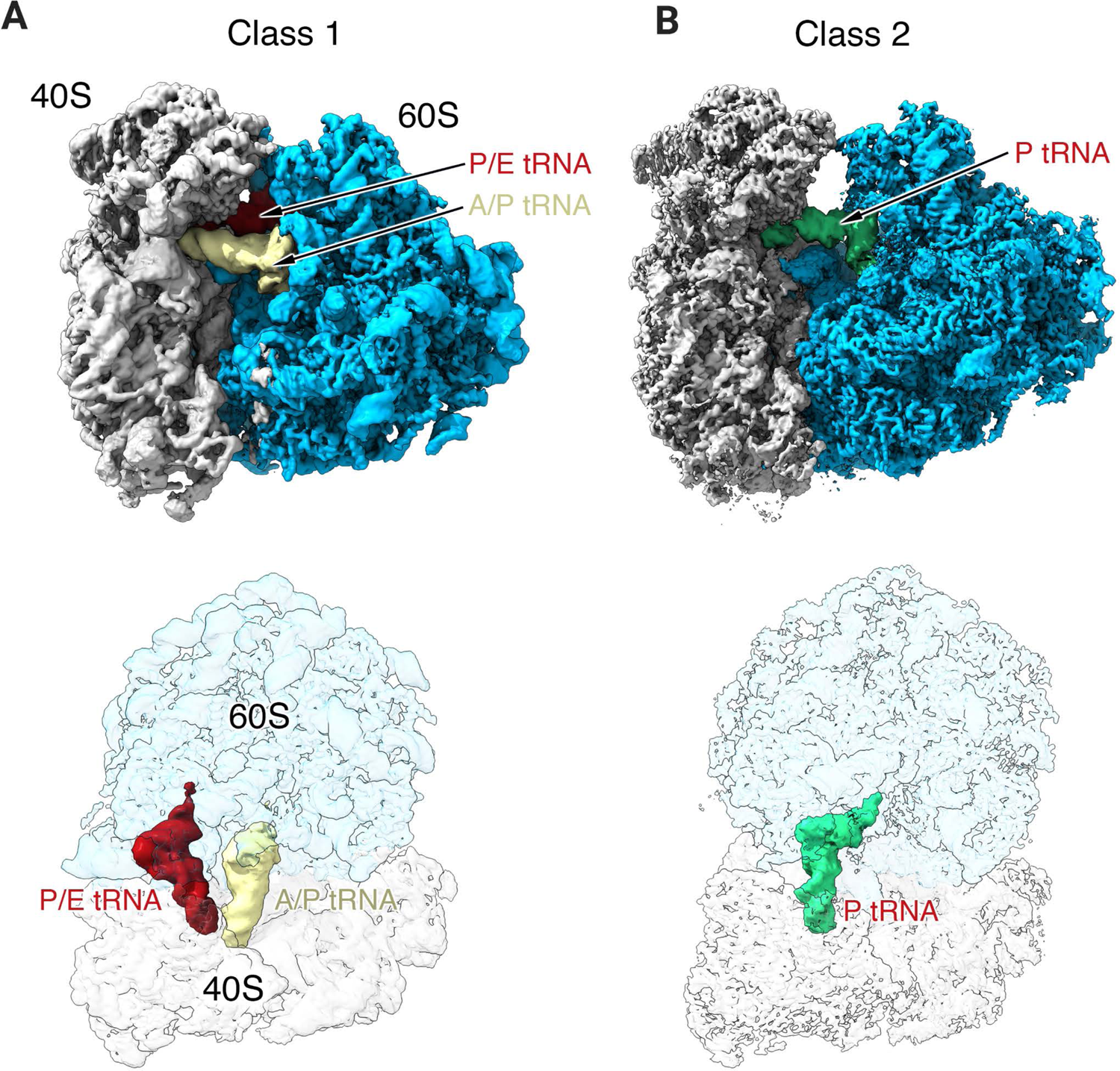
Cryo-EM of the monosomes digested from RNA granules. **(A)** Cryo-EM structure of the monosomes cleaved from RNA granules. (A) Cryo-EM map obtained for the major class of monosomes cleaved the RNA granules (Class 1). The top panel show a surface rendering representation where the tRNAs, large and small subunits are labeled. The bottom panel is a transparent view of the cryo-EM map view from the top. (B) Cryo-EM map for the minor class of ribosomal particles contained in the RNA granules (Class 2). This panel is organized as panel (A). The tRNA molecules loaded in each ribosome class are labeled.

These structures are consistent with ribosomes stalled in the elongation process and are similar to the structures seen in ribosomes stalled by collision, where the leading ribosome has an empty A site and the following ribosomes are stalled in the hybrid state (51). While it is not clear that these structures emanate from collided ribosomes (see discussion), the structures are consistent with the presence of stalled ribosomes in the pellet fraction. Despite the high resolution of these structures and the enrichment of FMRP and UPF1 in this fraction, no additional large densities were observed in these structures, although it is possible they were removed by the RNA digestion.

### Ribosome Footprints of the mRNAs protected by ribosomes in the granule fraction

After treatment with high concentrations of RNase I (1U), the cleaved ribosomes from the granule fraction were loaded onto a second 15-60% sucrose gradient and centrifuged to separate the monosomes before RNA extraction, library preparation, and sequencing of the footprint reads (Fig. 4A-C; 5A). UV absorbance (A254) shows a peak in fraction 2 (Fig. 4B) and EM of this fraction (which was used for profiling) confirms that it contains mainly 80S monosomes (Fig. 4C). As we are interested in stalled polysomes, no attempt was made to prevent ribosome run-off, and cycloheximide was not present during the tissue homogenization. We tested whether the presence or absence of cycloheximide impacts ribosome footprints (Supp. Fig 3). There was little effect of the omission of cycloheximide on the mRNAs detected by footprint reads as seen by a high Pearson’s correlation and clustering by principal components analysis (PCA) between relative numbers of footprint reads in the presence or absence of cycloheximide (Supp. Fig 3).

**Figure 4.**
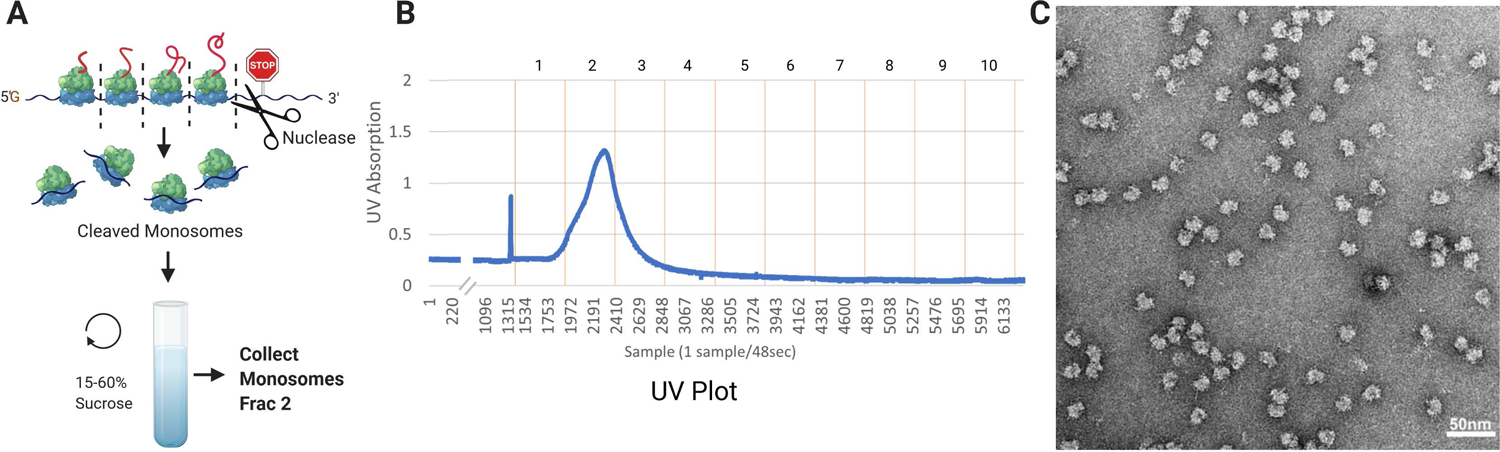
Purification of monosomes for ribosome footprinting. A. Schematic of the procedure, where the pellet is treated with nucleases and then sucrose gradient fractionation is used to isolate 80S monosomes. B) UV A254 absorbance of the sedimentation shows major peak in fraction #2. This is also the major fraction where S6 is found (Fig. 2F; Supp Fig. 1). C) Negative stained electron micrograph of Fraction 2 shows the presence of 80S monosomes in this fraction.

Similarly, nuclease digestion was performed after a brief treatment with high salt; while this treatment slightly improved the digestion (Fig. 2), the mRNAs detected by footprint reads were very similar to the salt untreated sample, again determined by a high Pearson’s correlation and clustering by PCA (Supp Fig 3). For consistency, only the samples prepared in the absence of cycloheximide and the presence of salt during digestion were used for the results reported below. Overall, we generated five biological replicates of libraries of footprint reads from the cleaved compacted ribosomes in the granule fraction (Fig. 5A). The results below are from averages of the five replicates after normalizing for transcript length and library size (39).

**Figure 5.**
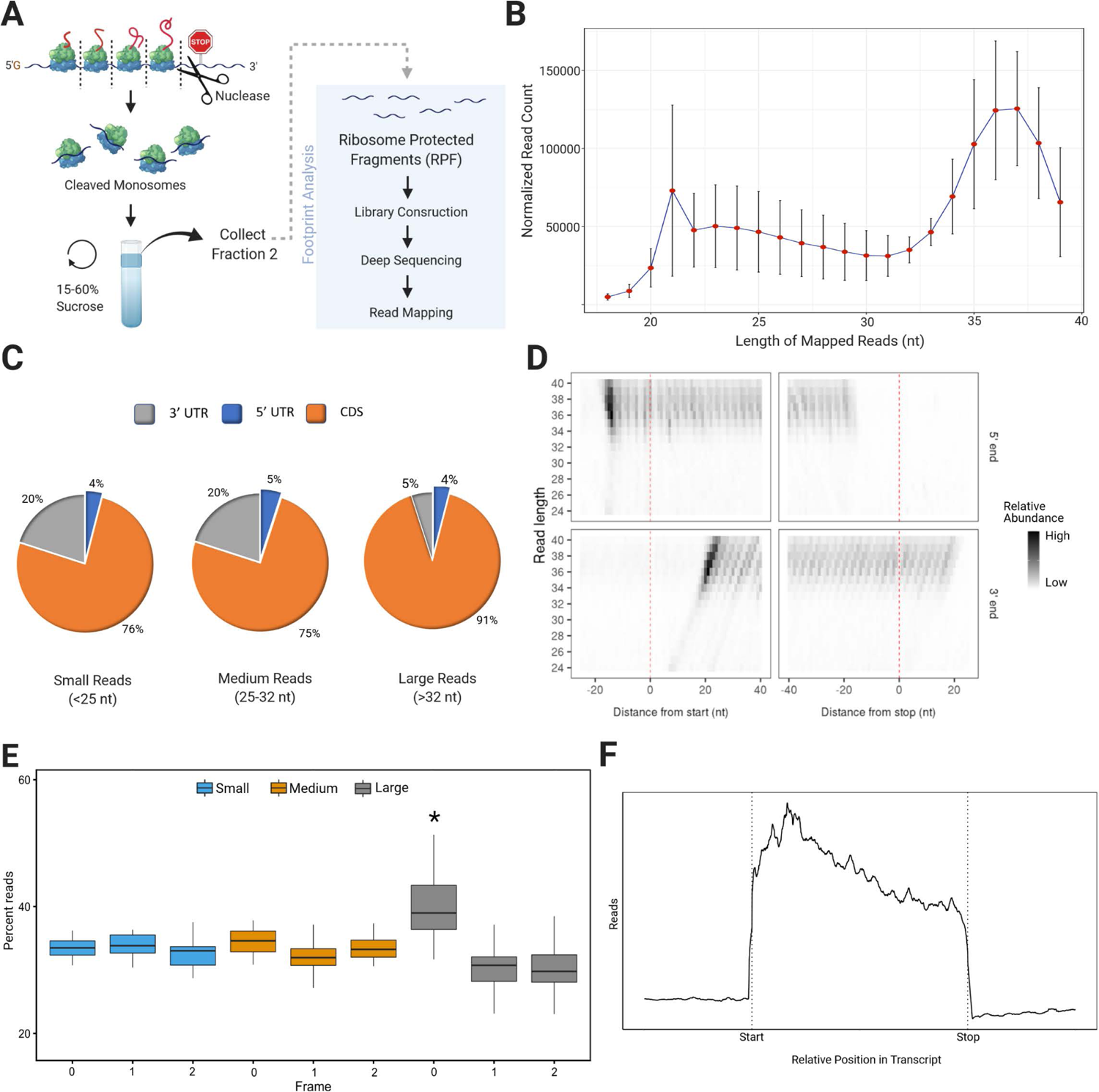
Ribosome footprinting of the granule fraction. (A) Diagram summarizing footprinting procedure. (B) Size distribution of normalized footprint reads, n=5 biological replicates, SEM. (C) Read coverage of different sized footprints (small <25 nt, medium 25-32 nt, large >32 nt) to 3’UTR, 5’UTR and CDS. (D) The number of read extremities (shading) for each read length (Y-axis) based on distance from start (left; 0 on x axis is A in ATG) and stop (right; 0 on x axis is last nucleotide of stop codon) with the beginning of the read (5’) on top and the end of the read (3’) on bottom. Data are shown for one biological replicate, but results are similar for all replicates. (E) Periodicity statistics indicate that long reads (33–40) in frame 0 have significantly more periodicity than frame 1 and frame 2 for long reads, or any frame for short (21–24) and medium reads (25–32) (ANOVA F (261, 8) =13.9, p<0.001; Post-Hoc Tukey HSD test, Long reads in 0 frame*, p<0.001 against long reads in frame 1 and frame 2 and other reads in frame 0) SD, n=39 long, 40 medium, 11 short (N is based on each read length in each biological replicate; not all read lengths are present in all biological replicates); Errors are SD. (F). Distribution of large reads with the CDS of all transcripts normalized to the same length shows that reads are biased to the first half of the transcripts. The X axis is the relative position in the transcript and the y axis is the average number of reads for that relative position. Data are shown for one biological replicate, but results are similar for all replicates.

Ribosome profiling reads are usually generated from canonical fragment sizes between 28 and 34 nucleotides (23). Surprisingly, the peak read size (36-37 nucleotides) from the ribosome protected reads was larger than this canonical ribosome protected fragment (Fig. 5A,B). It has been previously reported that classical ribosomes produce medium sized 27-29 nucleotide footprints and small 20-22 nucleotide footprints should the ribosome have an open A site (52). The larger reads we observed may be due to (i) an altered state of the stalled polysome, (ii) increased protection due to associated RNA binding proteins, or (iii) incomplete digestion. It is also possible that even larger reads are present since the libraries were generated after running a gel to size select the ribosome protected RNAs (see Methods). The longer reads map better to the coding sequence (CDS) than shorter reads (Fig. 5C). The presence of reads in the 3’UTR probably represents contamination from RBP complexes on the 3’UTR that co-migrate on the sucrose gradient with monosomes. Alignment at the start and stop codons showed that the excess length of the longer reads was due to extension at the 3’end of the footprint reads (Fig. 5D). While approximately 14 nucleotides were protected at the 5’ end, regardless of the read length, the extension at the 3’end increased with the read length giving a diagonal line on a plot of read length vs. distance from start or stop codon (Fig. 5D). Ribosome footprint reads should show periodicity due to the three-nucleotide code in the mRNA and reads over 32 nt showed higher periodicity than the shorter reads (Fig. 5E). We had predicted that footprint reads from stalled polysomes would be most enriched at the stop codon and the 3’ end of the message due to the requirement of UPF1 for the formation of stalled polysomes and the recruitment of UPF1 at the stop codon. However, we did not observe a bias for footprint reads near the stop codon. Instead, there was some bias in the large footprint reads for the first half of the message (Fig. 5F). Taken together, ribosome profiling of dissociated monosomes derived from the granule fraction from P5 rat brain reveals enrichment in large ribosomal footprints (>32 nt), which carry a prominent 3’end extension, display a preference for the first part of the mRNA CDS, and do not display a bias for the stop codon.

### Analysis of mRNAs that are abundant and enriched in footprint reads from the granule fraction

We were interested in identifying what mRNAs represent the most abundant constituent of the granule fraction and what mRNAs are enriched in this fraction compared to total mRNA levels. We calculated the average for Abundance (reads per kb of mRNA; RPKM) of footprint reads from the five biological replicates to determine the most abundant constituents of the granule fraction (Data set 1). To determine what mRNAs are enriched in these ribosomes, we calculated what is characteristically called the translation efficiency (abundance of ribosome footprints/abundance of total mRNA as determined by conventional RNA seq of the starting fraction) for each mRNA (Data Set 1). However, in the context of presumed stalled ribosomes, translation efficiency may be a misleading term, so we renamed this measure as enrichment for the present study.. Gene Ontology (GO) analysis of the 100 mRNAs with the largest abundance showed significant over-representation of mRNAs encoding cytoskeletal proteins (Table 1) that are expressed developmentally in neuronal projection and synaptic compartments (Fig. 6A), including Map1b (Table 1), an mRNA we had previously shown to be translated in dendrites through reactivation of stalled polysomes (8, 9). While cytoskeletal mRNAs dominate the most abundant GO category and are also enriched (Data Set 1), they are also relatively abundant in the total mRNA population. The GO analysis for enriched mRNAs showed a significant over-representation of mRNAs encoding RNA binding proteins or proteins involved in RNA metabolism (Fig. 6B), including the gene mutated in ALS; FUS (Table 2). These results were similar regardless of whether the 50, 200 or 500 most abundant or enriched mRNAs were selected for the analysis (Data Set 2,3) We next determined if mRNAs previously reported to be regulated by elongation or initiation in the nervous system were over-represented in our samples either through abundance or enrichment (Fig. 7). Strikingly, mRNAs whose translation is regulated by elongation through eEF2 phosphorylation (53) were significantly abundant and enriched in the footprint reads (Fig. 7A, B). In contrast, mRNAs regulated by initiation, through eIF4E phosphorylation (54), TOR activation (including TOP mRNAs) (55) or through upstream ORFs after eIF2 alpha phosphorylation was increased by the stimulation of metabotropic glutamate receptors (56), were not enriched in the preparation (Fig. 7A, B). While TOP mRNAs, mainly encoding ribosomal subunits are relatively abundant and known to be transported in neurons, we observed a significant de-enrichment of footprint reads from TOP mRNAs in footprint libraries of the compacted ribosomes (Fig. 7A), consistent with the notion that transported mRNAs blocked at initiation are not enriched in this preparation.

**Figure 6.**
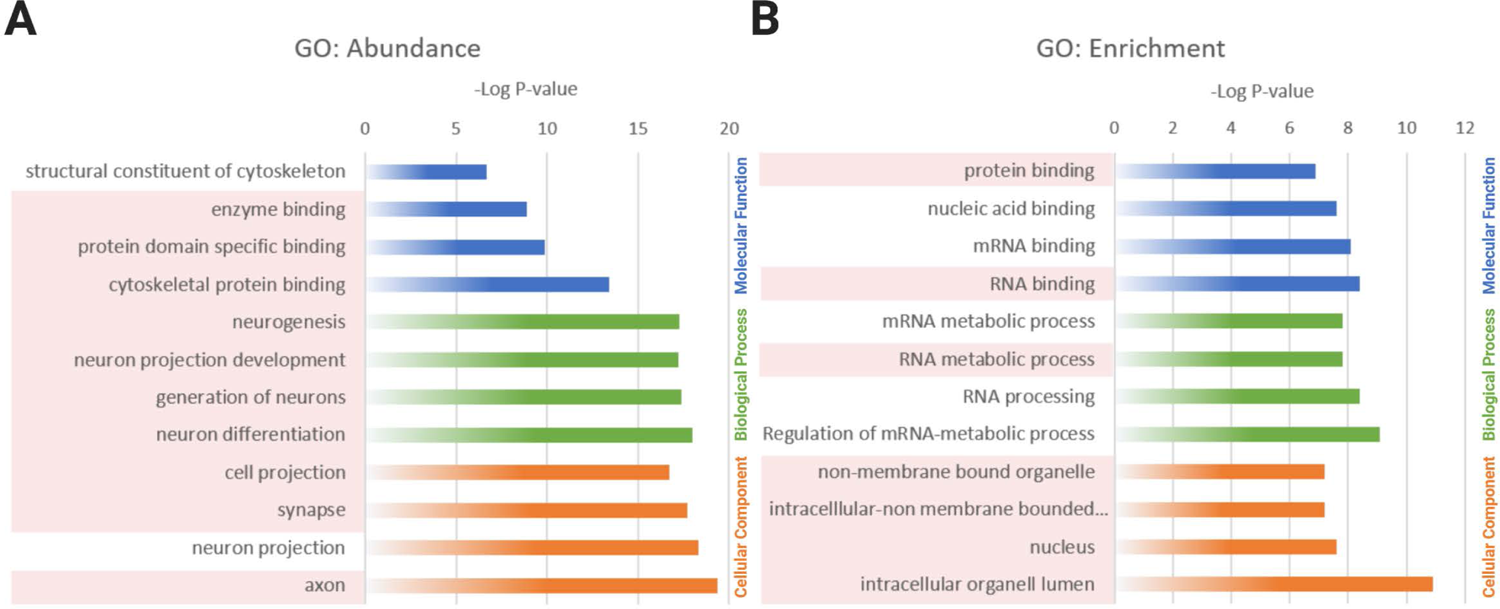
GO Analysis of footprint mRNAs. (A-B) Gene Ontology (GO) terms of selected comparisons for most abundant (A) and most enriched (B) mRNAs found in reads based on -Log P-values. Terms highlighted in red represent terms involving RNA binding for enrichment and cytoskeletal related terms for abundance. Complete table with all significant findings in Data Set 2.

**Figure 7.**
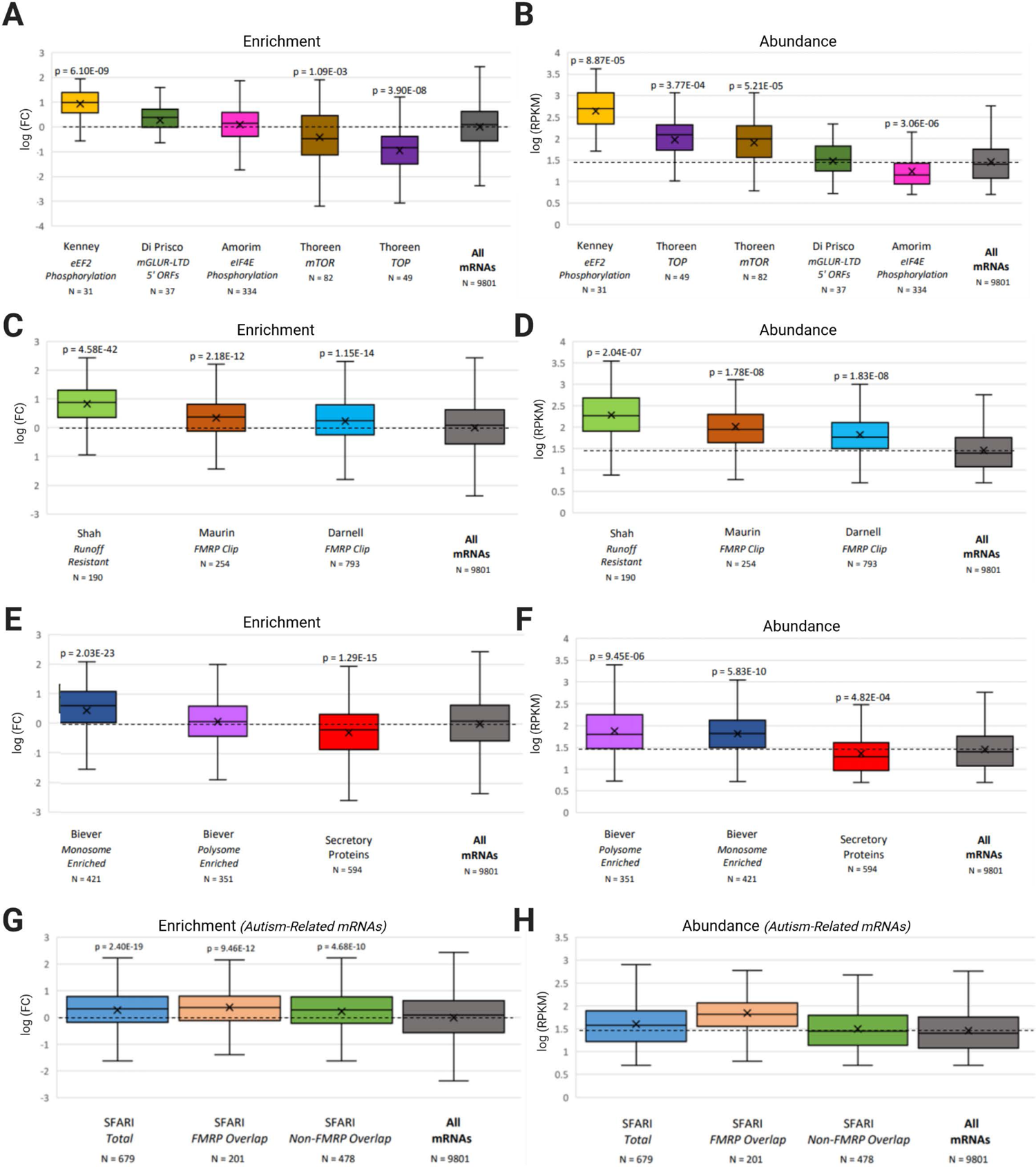
Correlation analysis of mRNAs abundant and enriched in ribosome footprints. (A) Enrichment and (B) Abundance comparison of footprint reads to mRNAs regulated by translation elongation (30) eIF4E phosphorylation (31) mTOR (32) TOP mRNAs (32) and mRNAs upregulated by mGluR with upstream open reading frames (33). (C) Enrichment and (D) Abundance comparison of runoff-resistant mRNAs (19) and mRNAs that are CLIPped by FMRP (17, 34). (E) Enrichment and (F) Abundance comparison of mRNAs translated preferentially by monosomal and polysomal ribosomes in the neuropil (35) and secretory mRNAs (secretory proteins with reviewed annotation from UNIPROT), compared to all mRNAs. (G) Enrichment and (H) Abundance comparison of autism-related mRNAs from the SFARI database (syndromic and levels 1-3). The total SFARI group was also divided into ones that are also in the FMRP CLIP group (17, 34). and ones that are not. For all groups there was a cut-off of 5 RPKM to avoid mRNAs not expressed in the nervous system. P values from comparison to all mRNAs (Students t test with Bonferroni correction for multiple tests (n=14 for all comparisons in figure). Only Significant P values (p<0.01 after correction are shown). The N for each comparison group is shown under the group.

**Table 1.** Top 20 most abundant mRNA. Top 20 most abundant mRNAs sorted on the Footprint reads per kilobase per million reads (RKPM) compared to Total RNA reads per kilobase per million reads (Total RNA RPKM). Also shown are the log fold change (Footprint RPKM compared to total RNA RPKM). Also shown are evaluations of the significance of the fold change (the P value and the False Discovery Rate (FDR). mRNAs highlighted in blue match the GO term for cytoskeleton constituent of cytoskeletal binding.

| Gene Name | logFC | P-value | FDR | Total RNA RPKM | Footprints RPKM |
| --- | --- | --- | --- | --- | --- |
| ACTB | 1.940985547 | 1.53E-08 | 9.23E-07 | 3311.57669 | 9408.327001 |
| TUBB5 | 1.033645345 | 1.37E-05 | 1.55E-04 | 3698.919325 | 7404.359382 |
| TUBB2B | 1.63257495 | 4.98E-07 | 1.13E-05 | 1913.684313 | 5570.961651 |
| YWHAE | 1.711033554 | 7.60E-09 | 5.65E-07 | 1688.064086 | 4751.885604 |
| TUBB3 | 1.244833653 | 2.40E-07 | 6.48E-06 | 2052.960915 | 4594.904687 |
| YWHAZ | 1.05722464 | 1.71E-06 | 2.96E-05 | 2004.358037 | 4216.85062 |
| STMN2 | 1.400937863 | 2.81E-06 | 4.44E-05 | 1692.26231 | 3719.643634 |
| CALM1 | 0.154694995 | 4.87E-01 | 6.19E-01 | 3093.888941 | 3625.962055 |
| DPYSL2 | 1.24010485 | 2.46E-09 | 2.57E-07 | 1568.286012 | 3524.363783 |
| YWHAG | 1.82197446 | 1.01E-09 | 1.32E-07 | 1166.611045 | 3340.758587 |
| CALM3 | -0.309008441 | 7.89E-02 | 1.58E-01 | 3403.092834 | 2994.609636 |
| CALM2 | 0.862970381 | 7.06E-04 | 3.78E-03 | 1549.807564 | 2953.038226 |
| UCHL1 | 1.003234991 | 4.34E-05 | 3.93E-04 | 1439.792879 | 2647.37674 |
| DPYSL3 | 2.223810027 | 1.16E-10 | 2.71E-08 | 698.2863694 | 2518.842074 |
| ZWINT | 0.887865185 | 8.91E-04 | 4.56E-03 | 1426.858532 | 2492.284205 |
| MAP1B | 1.865404024 | 1.12E-10 | 2.71E-08 | 755.4364087 | 2259.892443 |
| STMN1 | 0.273922938 | 2.08E-01 | 3.31E-01 | 1722.323579 | 2160.340529 |
| GAP43 | 0.569199253 | 1.27E-03 | 6.09E-03 | 1409.864902 | 2067.364803 |
| CRMP1 | 1.496025361 | 3.20E-09 | 3.00E-07 | 882.8247474 | 1959.479377 |
| YWHAB | 1.791487719 | 8.44E-09 | 6.04E-07 | 578.6707707 | 1808.787351 |

**Table 2.** Top 20 most enriched mRNAs. Top 20 enriched mRNAs sorted on the fold change (FC) in global footprint reads per kilobase pre million reads (Footprint RKPM) compared to Total RNA reads per kilobase per million reads (Total RNA RPKM). mRNAs were excluded if the RPKM was predominantly due to reads less than 32 bp (>2 fold) or if the total RPKM was less than 5. Also shown are two evaluations of the significance of the fold change, the P value, and the false discovery rate (FDR). mRNAs highlighted in red match the GO term mRNA or RNA binding.

| Gene Name | logFC | P-value | FDR | Total RNA RPKM | Footprints RPKM |
| --- | --- | --- | --- | --- | --- |
| FUS | 3.16434772 | 6.49E-10 | 9.56E-08 | 69.7831423 | 337.199324 |
| FAM168A | 2.94304308 | 7.81E-07 | 1.60E-05 | 52.1322914 | 130.2335745 |
| PRRT1 | 2.89433174 | 1.85E-06 | 3.16E-05 | 26.2478904 | 55.2638004 |
| NCL | 2.87065155 | 4.50E-13 | 4.21E-10 | 63.0689626 | 366.6690471 |
| RBM25L1 | 2.67673894 | 5.59E-08 | 2.27E-06 | 4.69685407 | 22.57140369 |
| HNRNPD | 2.63687085 | 5.04E-10 | 8.14E-08 | 135.136829 | 590.9433885 |
| GATAD2B | 2.55905455 | 1.39E-08 | 8.75E-07 | 34.7720695 | 118.9687961 |
| SOX9 | 2.45296112 | 1.01E-05 | 0.00012227 | 25.4556107 | 59.89774259 |
| UBQLN2 | 2.43529701 | 4.11E-11 | 1.25E-08 | 101.635634 | 385.068685 |
| HNRNPA2B1 | 2.43013679 | 5.74E-10 | 8.95E-08 | 158.368037 | 668.3078643 |
| PRR36 | 2.41730546 | 1.44E-06 | 2.58E-05 | 23.624002 | 46.51721335 |
| ZMIZ1 | 2.37502673 | 7.30E-07 | 1.53E-05 | 38.2509104 | 89.47685878 |
| LOC103694210 | 2.3707282 | 2.29E-07 | 6.30E-06 | 16.8011259 | 76.08347195 |
| TAF15 | 2.36234981 | 2.89E-07 | 7.61E-06 | 10.1106776 | 38.79032835 |
| MNT | 2.36190606 | 3.14E-08 | 1.55E-06 | 10.7221613 | 28.72871691 |
| AHCYL1 | 2.36154326 | 3.68E-08 | 1.70E-06 | 59.2481984 | 150.2720528 |
| HNRNPA3 | 2.35579919 | 1.63E-10 | 3.46E-08 | 53.2766435 | 197.1313486 |
| LOC108348151 | 2.35324232 | 1.93E-05 | 0.00020574 | 11.6881118 | 46.27995067 |
| VAMP2 | 2.30892138 | 4.34E-07 | 1.00E-05 | 473.446629 | 912.2714551 |
| SF3A2 | 2.29351137 | 3.99E-07 | 9.54E-06 | 20.2483045 | 41.1681424 |

We next determined whether mRNAs previously reported to be enriched in stalled polysomes are particularly enriched/abundant in the footprint reads of compacted ribosomes. Recently, a publication identified mRNAs protected by ribosomes resistant to ribosomal run-off in neuronal slices (19). While this publication focused on the proportion of each mRNA for which ribosomes ran-off, their data also identified the mRNAs with the most protected fragments remaining after a long period of run-off (60 min). The 200 runoff-resistant mRNAs with the most reads remaining at 60 minutes were significantly enriched and abundant in our preparation (Fig. 7C, D). FMRP, an RBP highly enriched in the sedimented pellet, has been associated with stalled polysomes (17, 18). The mRNAs associated with FMRP in neurons, identified through cross-linking immunoprecipitation (CLIP), are also more resistant to ribosome run-off (17). We examined the abundance and enrichment of two separate FMRP CLIP studies from brain tissue, and both were significantly enriched and abundant in our preparation (17, 57) (Fig. 7C, D).

We also examined several other datasets to evaluate the mRNAs with footprint reads in the pelleted compact ribosomes. Secreted mRNAs are stalled by their signal peptide and then co-translationally inserted into the endoplasmic reticulum (ER). The transport of secreted mRNAs stalled at elongation would also involve the transport of ER in the RNA granules; while transport of ER is possible, we suspected that secretory mRNAs would be de-enriched in the granule fraction, and this was indeed the case (Fig. 7E). Recently, mRNAs that are preferentially translated from monosomes in neuronal processes were identified (36). While we predicted that these mRNAs would also be depleted from our preparation, they were significantly enriched (Fig. 7E), while mRNAs preferentially transported in polysomes were not significantly enriched (Fig. 7E), although both types of mRNA were abundant in our preparation (Fig. 7F). Total mRNA levels for the preferentially polysomal transported mRNAs were significantly higher than the total mRNA levels for the preferentially monosomal translated mRNAs at this developmental timepoint (158 ± 17 polysome RPKM (n=327) vs. 72 ± 4 monosome RPKM (n=458), S.E.M, p<0.001 Student’s t-test).

Since translation from stalled polysomes is implicated in neurodevelopmental disorders, we examined if protected reads from Autism-related genes from the Simons Foundation Autism Research Initiative (SFARI) database were enriched and abundant in the protected reads. Compared to all mRNAs, these mRNAs were significantly enriched in our data (Fig. 7G, H). There is a significant overlap with FMRP CLIPped mRNAs in this dataset, but both FMRP CLIPped SFARI mRNAs and non-CLIPped SFARI mRNAs were equally enriched in the protected reads (Fig. 7G, H).

Finally, some of the enrichments we observed may be due to our preparation being specifically from the nervous system. To account for this bias, we repeated this analysis using only mRNAs known to be transported in neuronal processes (36). While the mRNAs known to be transported in neuronal processes were significantly enriched and abundant in our preparation (Supp Fig 4) when we restricted the total set of mRNAs to only include this set, all of the results above were replicated (Supp Fig 5), suggesting our results are not biased due to enrichment of neuronally transported mRNAs.

### The ribosome protected reads are enriched in sequences matching FMRP CLIPs

If stalled polysomes are indeed enriched in the compacted ribosome-containing pellet, the footprint reads should help identify where on the mRNA ribosomes are stalled. Examination of the distribution of the large reads (>32 nt) on individual messages revealed highly non-uniform distribution of reads on mRNAs (Fig. 8A). As peaks in ribosome protected fragments are often non-reproducible (58), we used stringent criteria to identify peaks. First, peaks were defined for each mRNA in each library based on a maximum value higher than the average RPKM for the mRNA and a minimum width of 18 nucleotides at the half maximum height. Second, the peak had to be present in at least 3 of the 5 biological replicates (Fig. 6B; Supp Fig. 5). Using these stringent criteria, we identified 766 peaks in 524 mRNAs (Data Set 4). While 90% of the mRNAs had only one or two peaks, Map1B, the mRNA most associated with stalled polysomes, had the largest number of identified peaks (10; Fig. 8A).

**Figure 8.**
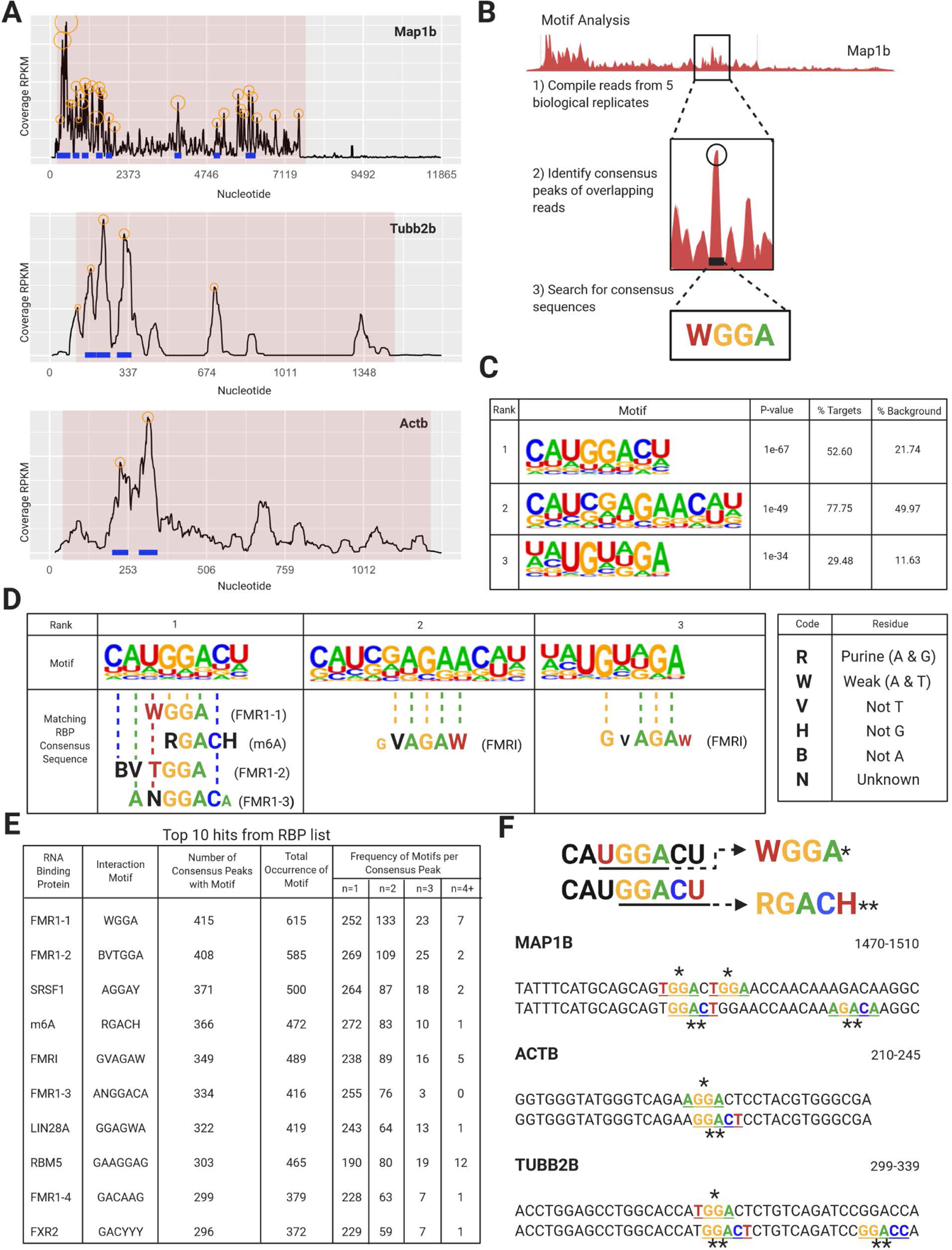
Sequences underlying ribosome protected fragments are enriched in sequences matching FMRP CliPs. (A) mRNA profiles of Map1b, Beta-actin and Tubulin 2b showing reproducible consensus peaks in the CDS; circles represent consensus peaks of footprint reads mapping to the same sequence, blue lines represent reproducible consensus peaks across biological replicates, red shading represents CDS. All replicates are shown in Supp. Fig 6. (B) Diagram summarizing how motif analysis is done. (C) Results from the HOMER program show the only 3 consensus sequences above the cut-off provided by HOMER. (D) Homer identified motifs overlapped with matching interaction motifs for RBPs listed in brackets. Residues that do not match are given in smaller font. RIGHT: Code key for residue annotation. (E) Table of top 10 RBPs with RNA interaction motifs present in consensus peaks. The number of peaks with multiple hits is also shown as Frequency of Motifs per Consensus Peak; n=x represents the number of motifs per consensus peak. All RBPs motifs examined are shown in Supp. Table 4. (F) Top ranked consensus sequence from HOMER showing overlapping sites for interaction motifs WGGA and RGACH, and their corresponding residues on a single peak from Map1b, Beta-actin and Tubulin 2b. Both sequences given for each protein are identical, but they have been annotated to show clusters of motifs that map to the same consensus sequence. Numbers on the right indicate the location of the sequence in each mRNA that correspond to a consensus peak (blue) on the mRNA profiles shown in (A).

Contrary to our initial hypothesis that peaks representing stalled footprint reads would be clustered around the stop codon, only 6 of the 766 total peaks were at the stop codon, and, similar to the overall coverage of footprint reads, the peaks were biased to the first half of the message with an average position of 0.37 ± 0.26 (SD) where 0 is the start codon, and 1 is the stop codon (Data set 4). The average length of the sequence within a peak was 36 nt ± 6 nt, similar to the most common footprint read size (Fig. 5B), and there was no clear evidence for the presence of collided ribosomes (peaks of twice the size of a ribosome protected fragments). We also identified peaks in the smaller reads, but only 65% of these peaks were in the CDS, and there was little overlap with the peaks in the long reads. In contrast, 94% of the peaks from large reads were in the CDS.

We next examined whether consensus sequences were in these peaks of footprint reads from the large fragments. We used an unbiased sequence motif search approach with the HOMER program (46). HOMER uses relative enrichment and requires a background sequence. To remove usage bias for mRNA sequences, we used similarly sized fragments from mRNAs with no peaks as our background selection. The Homer program identified three highly significant consensus sequences in the peaks above the false discovery rate determined by the program (Fig. 8C). Notably, the most significant consensus sequence (p=1e-67) included the consensus sequence (WGGA) previously derived from analyzing FMRP CLIP sequences (20, 21), which also overlapped with the consensus sites for m6A methylation (RGACH or RRACT) in the nervous system (24) (Fig. 8D-F). The motif was not biased to the start or end of the protected reads (average position of the motif in reads was 19 ± 9 bases) and thus did not represent sequences selected because they were difficult for nucleases to digest, since sequences selected due to resistance to digestion would be at the end or beginning of the read. Analysis of 36 nt in front of the peaks or 36 nt behind the peaks did not result in a HOMER consensus sequence above the false discovery rate determined by the program. Similarly, peaks identified using small or medium-sized reads also did not result in a HOMER consensus sequence above this cut-off.

We also performed a directed search of the peaks for sequences matching consensus binding sites of RBPs (59) using FIMO (44). Consistent with the non-biased search, FMRP CLIP consensus sites had the most matches (Fig. 8D, E; Data Set 5). Strikingly, these motifs mainly recognized purine-rich sequences (Fig. 8E; 76% of nucleotides in the top 10 RBP motifs are purines). However, the overall peak nucleotides are not enriched in purines compared to the background sequences (both at 57% purines). The abundance of purines in the consensus peaks suggests a possible role for the RBP PURA, a protein highly enriched in the granule fraction, but without a known consensus binding site. Of the 766 peaks, there were 415 peaks with a WGGA site and also many peaks with multiple WGGA sites in the peak making a total of 615 WGGA sites in consensus peaks (Fig. 8E; Data Set 4). The number of multiple matches (39% of peaks with a WGGA have multiple WGGA sites) was more than those found in the corresponding background sequences (11 ± 1 %, from 10 separate selections of 750 background sequences with a WGGA peak). Of the 615 WGGA sites found in the peaks, 182 WGGA sites overlapped with an m6A site (examples shown in Fig. 8F). This percentage of WGGA sites overlapping with an RGACH site (30%) was significantly more than the number of overlaps seen in WGGA sites in our background samples (19 + 1.3%; 10 separate selections of 1300 WGGA sites from background sequences). Moreover, there were also many examples where m6A sites and WGGA sites were in the same peak but did not overlap. Of the 415 peaks with a WGGA site, 315 (76%) had an m6A consensus site in the same peak significantly more than were found in the background sample (39 ± 4%, 10 separate selections of 750 background sequences with a WGGA peak).

There were also many matches to other FMRP CLIP consensus sites (Fig. 8D). After including GVAGAW and GACAAG, over 80% of the 766 consensus peaks contained an FMRP CLIP sequences or m6A consensus sites suggesting a strong sequence bias for these sites in the regions of the mRNA enriched in footprint reads (Data Set 4).

## Discussion

### The granule fraction is an enriched preparation for stalled ribosomes

Previous reports showed that increased density could be used to segregate RNA granules from normal polysomes (7,12–14). However, these studies did not show that ribosomes in these fractions are stalled. An association between stalled polysomes and compacted ribosomes was first made based on FMRP association (17) however, the RNA granules were not isolated by sedimentation in that study. Here, we show that the sedimented pellet (granule fraction) containing clusters of ribosomes is not only enriched for FMRP (as had been previously shown (13)), but also for UPF1, an additional protein functionally implicated in stalled polysomes (8). Moreover, mRNAs that were previously shown to resist ribosomal run-off by initiation inhibitors (19), or to be regulated at the elongation stage (53) were over-represented in the footprint reads generated from this fraction. Finally, cryo-EM of the monosome cleaved from the ribosome clusters in the GF demonstrate that they are stalled. These findings are consistent with the granule fraction enriching for ribosomes that were stalled in neurons.

The fraction is unlikely to be just larger polysomes. First, there are very few ribosomes in the fractions preceding the pellet, inconsistent with a gradual running off of large polysomes. Second, there is no correlation between the abundance or enrichment of mRNAs in the GF based on their length (total transcript or ORF) (Supp. Fig. 7), as would be expected if these were simply larger polysomes. Third, counting ribosomes in the cluster did not detect larger polysomes in the pellet compared to the polysome fraction. The pellet/GF may represent aggregates of ribosomes/polysomes. However, the question would remain why these would show specific enrichment of both specific types of mRNA (particularly those known to be stalled in neurons) and proteins (FMRP, UPF1). It is possible that stalled polysomes or RNA granules containing stalled polysomes are more likely to aggregate than normal polysomes.

It should be noted that this preparation represents a snapshot of P5 brains. This sedimentation protocol is complicated by the presence of myelin at later stages of development (13). Still, if feasible, the mRNAs found in the granule fraction would likely be different at the adult stage since many of mRNAs with the most abundant footprint reads that we saw in this fraction are developmentally implicated in neuronal outgrowth. A related protocol developed for E18 rat brains showed fewer ribosomes when performed in adults (14). However, stalled polysomes and some of the mRNAs isolated here (notably Map1b) are implicated in mGluR-LTD, a plasticity present mainly in mature brains (60). Thus, it is plausible that the role of stalled polysomes, or the mRNAS that they regulate, may change as the brain develops.

### Cryo-EM analysis reveals two populations of stalled ribosomes

The structure of these ribosomes very much resembled those of the previously described “collided” ribosomes (51). Collided ribosomes appear when a trailing ribosome encounters a slower or “paused” leading ribosome in a polysome. The “paused” leading ribosome contains a peptidyl-tRNA in the P-site, and the trailing collided ribosome adopts the rotated state with A/P and P/E hybrid tRNAs. However, we did not observe other signs of collided ribosomes, such as periodic peaks or a size of peaks corresponding to two or more ribosomes. Indeed, ribosomes that are stalled due to collision with a downstream ribosome recruit specific factors to resolve the stall and rescue the ribosomes (61). These factors are not present in proteomics of RNA granules (13–15), including proteomics from a very similar protocol (13). Indeed, if these stalled ribosomes are meant to be transported and reactivated, they need to be protected from the surveillance mechanisms that normally target prematurely stopped or collided ribosomes, which leads to unstalling and/or degradation of the proteins (61).

15% of the monosome structures had an empty A site. It is unlikely that the large protected fragments are due to these structures as only 15% of the monosomes has this structure, but the majority of protected reads are large. Moreover, ribosomes with unoccupied A sites should give smaller protected fragments (52). Thus, the large reads and the peaks of protected fragments are presumably coming from the ribosomes blocked in the hybrid position or prior to eEF2 mediated translocation. Again, this differs from what would be expected from collided ribosomes where the ‘paused’ ribosome has an empty A site, while the following ribosomes are stalled in the hybrid position.

We found no additional densities that could be assigned to putative stalling factors at the tRNA interface region of the ribosome or other regions of the ribosome. This is despite the enrichment of FMRP in this fraction and its proposed role as a ribosome binding stalling factor. Thus, these structures are more consistent with stalling being encoded by the consensus mRNA sequences instead of a specific protein factor. However, more focused examination will be required to identify small molecules, post-translational modifications, or other mechanisms that may underlie the stalling mechanism. It is also possible that nuclease treatment removed stalling factors located on the mRNA outside the protected fragments, although we did not find consensus sequences immediately before or after the peaks of protected fragments. Further cryo-EM studies on the clusters before nuclease treatment may be able to address this issue and perhaps provide more insight into the unexpected size of the protected fragments.

### Cytoskeletal and RNA binding proteins represent abundant and enriched mRNAs, respectively

Axon and dendrite outgrowth, coupled with synapse formation, occur at high rates in P5 rat brains (62). Indeed, the most abundant footprint reads are on cytoskeletal mRNAs and mRNAs encoding proteins that are highly enriched in growth cones (such as 14-3-3 proteins)(63). The most abundant number of footprint reads is found on beta-actin, whose local translation in both axons and dendrites is important for neuronal outgrowth (64, 65). Somewhat more surprising was the high enrichment for RNA binding proteins in the footprint reads (Fig. 6; Table 2). This suggests an important homeostatic aspect for translation from stalled polysomes, where the increased translation of RBPs will have critical effects on the translation of other messages not necessarily present in stalled polysomes.

### Identification of conserved motifs enriched in footprint read peaks from compacted ribosomes

The finding that the footprint reads derived from the granule fraction are distributed mainly in large peaks is consistent with the enrichment of stalled ribosomes in this fraction. These peaks are strikingly enhanced in sequences previously defined as enriched in FMRP CLIPs (Fig. 7). This is consistent with the strong enrichment of FMRP in the granule fraction (Fig. 1) and the abundance and enrichment of mRNAs previously identified as associated with FMRP using CLIP experiments (Fig. 6). While most FMRP CLIP consensus sequences have not been directly shown to bind to FMRP, the WGGA sequence can be directly bound by FMRP (21). However, the major FMRP binding sites (G quadruplex and Kissing sequence) are not enriched in FMRP CLIP consensus sites (20). Moreover, if ribosomes protect this sequence, it is unclear how FMRP would gain access to these sequences. However, it is possible that FMRP initially binds this sequence, and this is followed by ribosome occupation and stalling. It is also possible that these sequences are enriched in FMRP CLIPs because FMRP is specifically associated with stalled ribosomes and these sequences specify where ribosomes would be stalled independently of FMRP. In this scenario, the sequences would not be directly bound by FMRP. Instead, FMRP would be crosslinked to sequences near to, but not protected by the ribosome. Since the CLIP sequences are approximately 100 bp, this is entirely consistent with both the CLIP and ribosome footprint read data.

GGA was also identified as a consensus site in the 3’UTRs of mRNAs whose transport to distal sites in neurons was decreased in the absence of FMRP(66). However, in this case, the GGA sites were involved in forming G quadruplexes and binding to the RGG domain of FMRP, while the WGGA CLIP sites in the open reading frame are not associated with G quadruplexes (20). Moreover, the lack of transport was largely rescued by the I304N mutant of FMRP that does not bind ribosomes (66). While this manuscript compared ribosome footprints of a cell line differentiated into a neuron-like cell in the presence or absence of FMRP, there was no indication that footprints from RNA granules were measured since the average size of the protected fragments examined were the standard 28-32 bp long.

The consensus sites in the peaks are also enriched in a consensus site for m6A modification. Interestingly, mRNAs with m6A sites are selectively transported in neurons (67) and this methylation plays an important role in neurodevelopment (68). Moreover, mRNAs that are associated with FMRP by CLIP experiments have previously been shown to be highly enriched for m6A modifications in neurons (24) and this has been proposed to play a role in FMRP-mediated nuclear export (69, 70). However, whether FMRP directly binds to m6A, interacts with m6A readers, or is associated with m6A through some other indirect interaction is unclear. There has been some indication that m6A directly leads to the stalling of ribosomes (71) or it may be that some m6A reader in neurons is important for the stall. Again, it is unclear how the reader would access the mRNA sequences protected by the ribosome, but similar to FMRP the sequence may be recognized first and then later occupied by the ribosome. It will be interesting in the future to determine the specific relationship between m6A methylation and stalled polysomes and whether initial findings of specific roles for m6A methylation in the developing brain are linked to their possible role in stalling translation in RNA granules.

### Stalled Polysomes or Stalled ribosomes?

We did not search for disome or higher periodicity ribosome footprints (72), as we performed ribosome footprints on purified monosomes. Nevertheless, we do not find any data suggesting that ribosomes are collided. Our data is, instead, consistent with a model in which a controlled form of stalling attracts specific factors in neurons, such as FMRP, that likely play a role in the compaction of the stalled ribosome/polysome and package the structure into a granule for transport before collisions occur. Thus, the ribosomes in the GF may not represent collided ribosomes on a polysome, but collections of ribosomes from distinct mRNAs packaged together. Indeed, the mRNAs preferentially translated by monosomes in dendrites are enriched in this compacted ribosome pellet. The packaging of many distinct mRNAs in the same mRNA granule is consistent with the finding that *in situ* RNA granules containing stalled polysomes probably contain many distinct mRNAs (73).

## Conclusions

While there have been assumed links between the ribosomes that sediment in sucrose gradients, stalled polysomes identified by resistance to ribosomal run-off in neuronal dendrites and FMRP’s association with stalled polysomes, there were previously no direct connections between these various lines of research. Identifying an enrichment for FMRP CLIP consensus sequences in protected reads in ribosomes from the pellet of sucrose gradients establishes these links. Notably, most investigations of translation regulation in neuronal tissues do not consider the pellet fraction after separating polysomes using sucrose gradients. Thus, they are not accounting for this pool of translationally repressed mRNAs. For ribosome profiling from the initial polysome pellet, the use of translation efficiency needs to be re-evaluated because many of the ribosomes in this initial pellet are presumably stalled. Moreover, some studies only include footprints of the canonical size and may exclude the larger size footprints that we observed in this preparation. Thus, these results have large implications for the interpretation of many studies on neuronal translation.

Our data strongly supports a model in which mRNAs of cytoskeletal and RNA binding proteins important for neurodevelopment are regulated through ribosomes stalled in elongation that are packaged into RNA granules and transported to distal sites where they, upon stimulation, would be reactivated to result in fast and local protein synthesis. RNA granule proteins such as FMRP, that interact with these mRNAs either directly by binding to stalling sequences on the mRNA or to stalled ribosomes occupying these sequences, would thus act as master regulators of the fate of many mRNAs through this novel mechanism of initiation-inhibitor resistant translational control.

## Supporting information

data sets 1-5

## Acknowledgements

We thank staff at the Facility for Electron Microscopy Research (FEMR) at McGill University. FEMR is supported by the Canadian Foundation for Innovation, the Quebec government, and McGill University. We thank Mehdi Amiri and the Sonenberg lab for their help with the acquisition of the UV absorption data. Figures were created using Biorender.com

## Data Availability

All raw reads from these experiments have been deposited at the GEO dataset at NCBI (GEO accession GSE173464). All excel files with complete numbers are attached as supplemental files. The cryo-EM maps obtained in this study have been deposited in the Electron Microscopy Data Bank (EMDB), and the accession codes are detailed in Supplementary Table S1.

## Supplemental Figure Legends

**Supplemental Figure 1.**
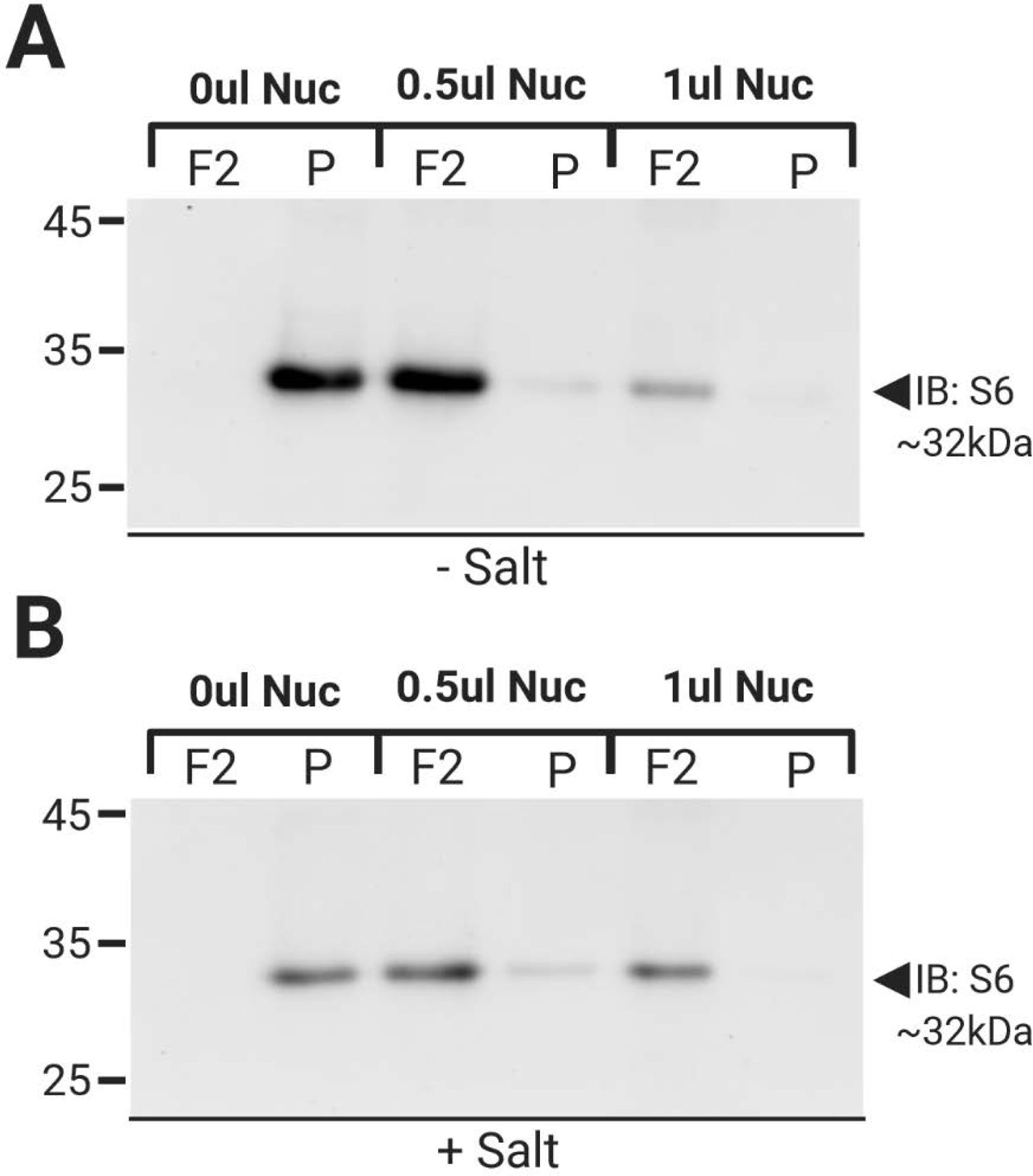
Complete digestion of pellet fraction. Western Blot analysis for S6 ribosomal protein of purified fractions collected after first treating the pellet from the first sucrose gradient with 0ul, 0.5ul and 1ul RNase I, with or without pre-treatment of 400 mM NaCl (-SALT: A; +SALT: B), followed by a second sucrose gradient (see Methods). Only fraction 2 was loaded in this experiment. In this example, almost complete digestion was observed even with lower nuclease concentrations and no salt. Increased nuclease digestion appeared to be related to different batches of nuclease and different amounts of time nuclease was stored, but this was not systematically analyzed.

**Supplemental Figure 2.**
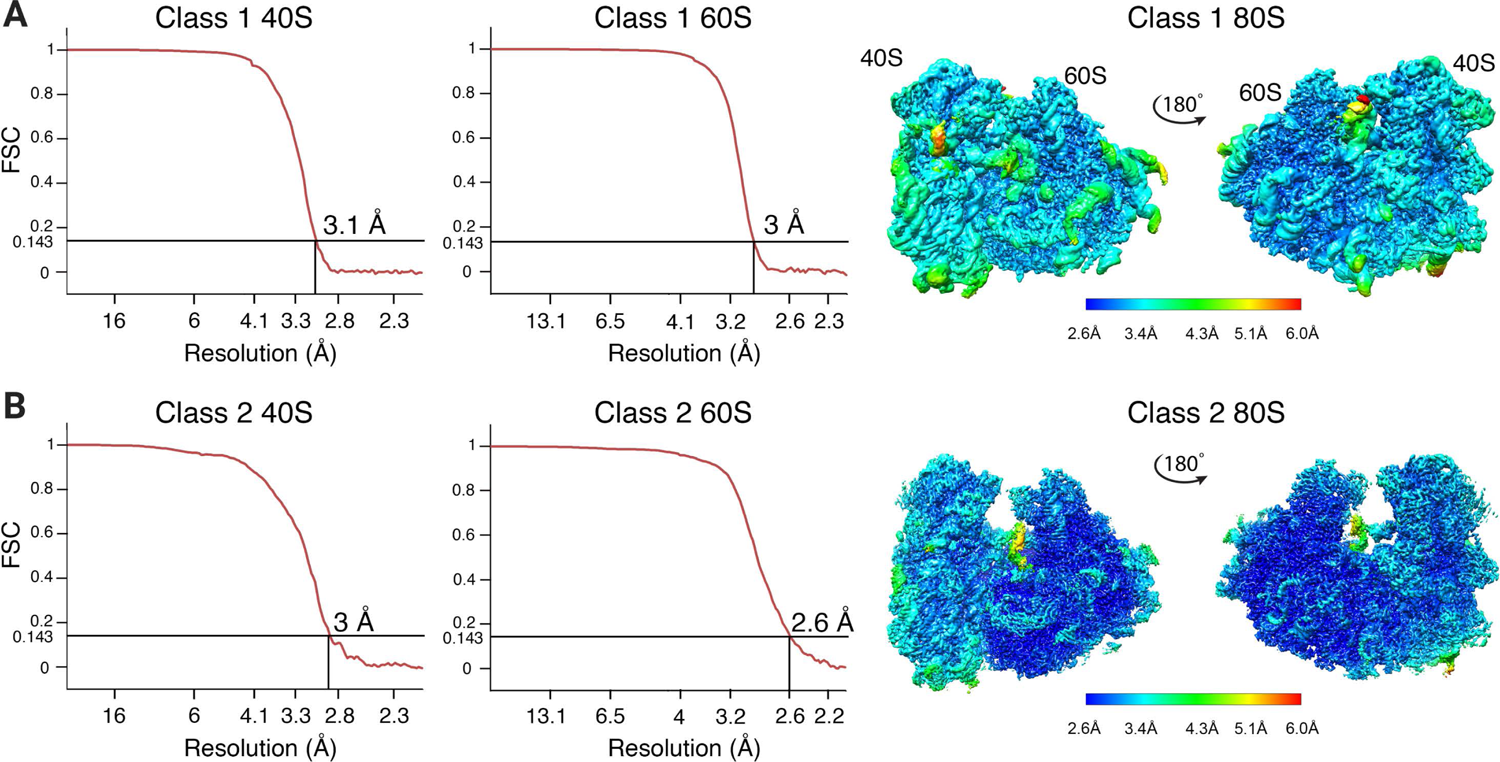
Resolution analysis of the two major classes of monosomes cleaved from the RNA granules. The cryo-EM maps for class 1 (A) and class 2 (B) of monosomes cleaved from the RNA granules were refined by multi-body refinement by dividing the 80S ribosome into two major bodies, the 40S and the 60S particles. Because each subunit was refined independently, we show the Fourier shell correlation graphs (left and middle panels) for each class. Graph are labeled to indicate whether they correspond to either the 40S or 60S subunit part of the cryo-EM maps. We used a FSC threshold of 0.143 to report the resolution. The right side of each panel shows the local resolution analysis of the cryo-EM maps obtained for both classes. These maps were obtained by merging the 60S and 40S cryo-EM maps obtained by multi-body refinement using the command ‘vop add’ in Chimera. Maps are colored according to their local resolution using the color coding indicated in the scale bars. Main structural landmarks are indicated.

**Supplemental Figure 3.**
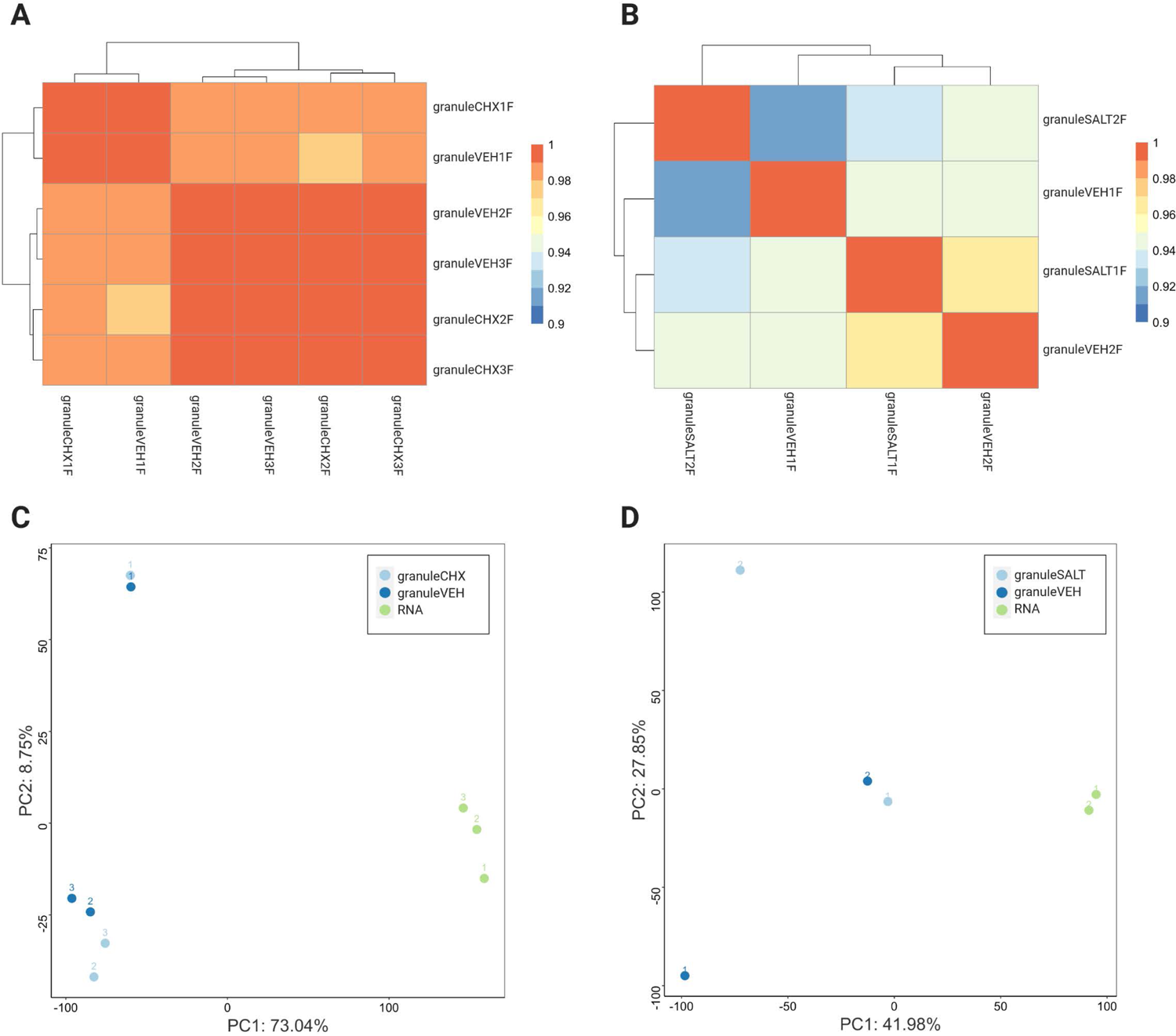
No effect of Salt or Chx treatment on RNA reads. (A-B) Heat map for the comparison of biological replicates accomplished in the presence or absence of cycloheximide (Chx) (A) or for the comparison of biological replicates accomplished in the presence or absence of salt (B). Warmer colors indicate a higher correlation between groups. Differences between biological samples were equal to or higher than the differences seen with treatment (C-D) Principal component analysis for the comparison of biological replicates accomplished in the presence or absence of cycloheximide (Chx) including RNA-SEQ of starting material (C) or for the comparison of biological replicates accomplished in the presence or absence of salt including RNA-SEQ of starting material (D). Differences between biological samples were equal to or higher than the differences seen with treatment. The footprint reads were clustered separately from the RNA-SEQ of total mRNA. All calculations were done with the Limma program (28).

**Supplemental Figure 4.**
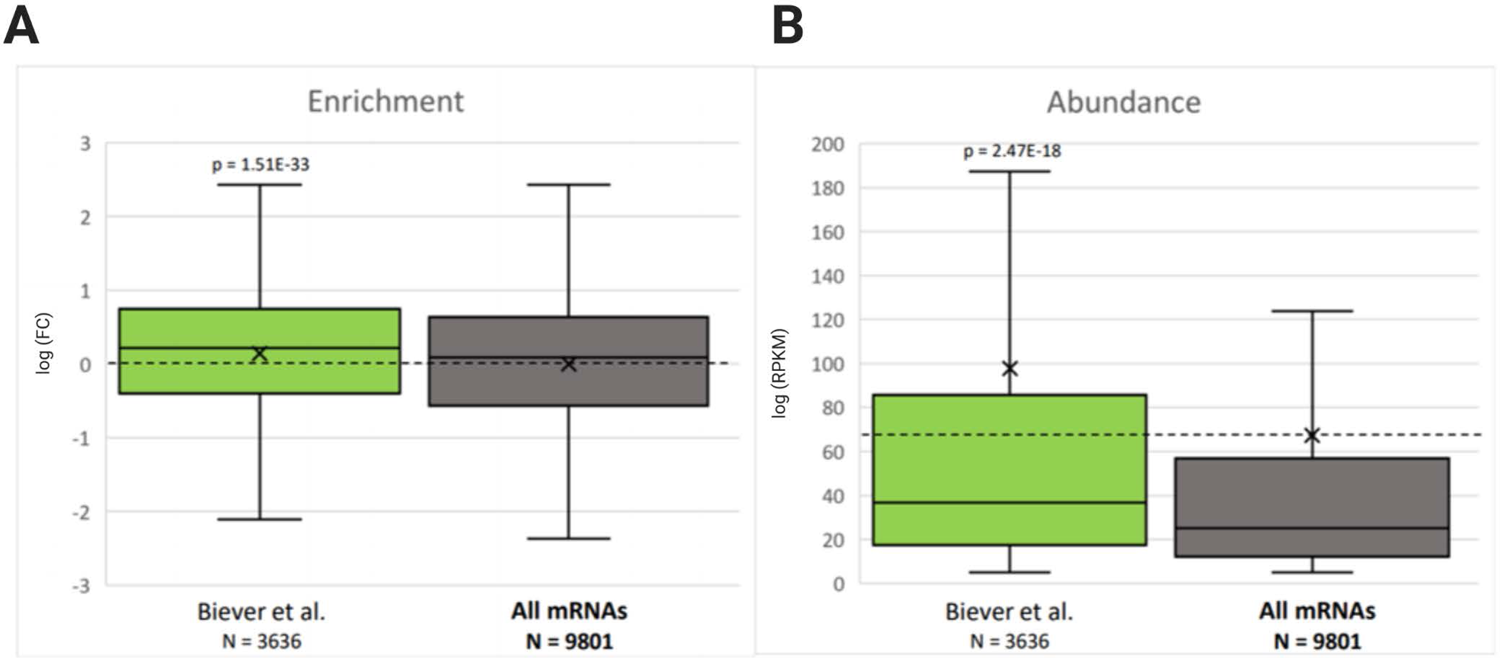
Enrichment and Abundance compared to transported mRNAs. (A) Enrichment and (B) Abundance comparison of footprint reads to mRNAs classified as transported to distal sites as determined (35). For all groups, there was a cut-off of 5 RPKM to avoid mRNAs not expressed in the nervous system. P values from comparison to all mRNAs (Students t-test with Bonferroni correction for multiple tests (n=2 for all comparisons in figure). Only Significant P values (p<0.01 after correction are shown). The N for each comparison group is shown under the group.

**Supplemental Figure 5.**
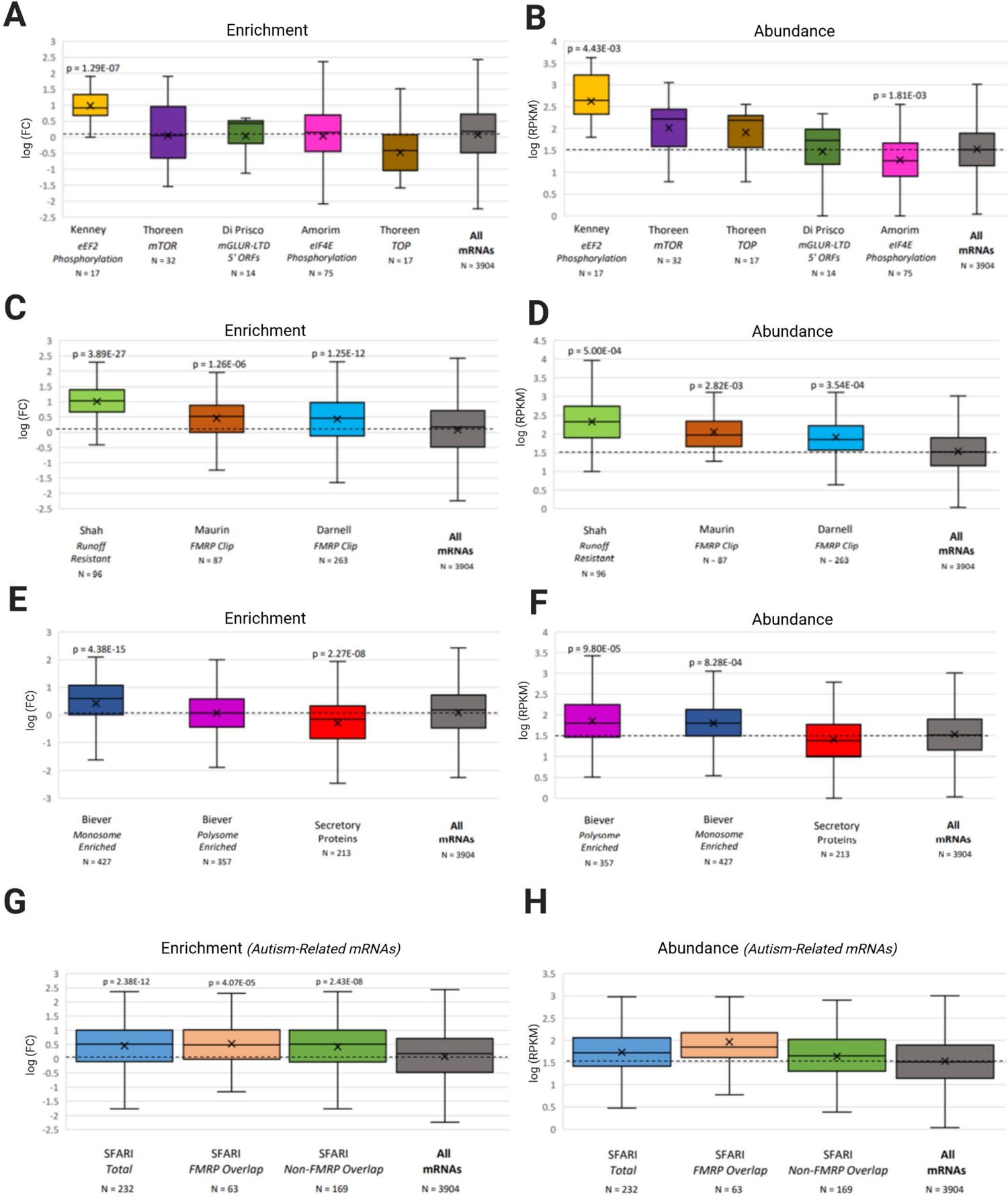
Comparisons only using neuronal transported mRNAs. (A) Enrichment and (B) Abundance comparison of footprint reads of all transported mRNAs (35) to transported mRNAs regulated by translation elongation (30) eIF4E phosphorylation (31) mTOR (32) TOP mRNAs (32) and mRNAs upregulated by mGluR with upstream open reading frames (33). (C) Enrichment and (D) Abundance comparison of all transported mRNAs to runoff-resistant mRNAs (19) and mRNAs that are CLIPped by FMRP (17, 34). (E) Enrichment and (F) Abundance comparison of all transported mRNAs to mRNAs translated preferentially by monosomal and polysomal mRNAs in the neuropil (35) and secretory mRNAs (secretory proteins with reviewed annotation from UNIPROT). (G-H) Enrichment and Abundance comparison of autism-related transported mRNAs from the SFARI database (syndromic and levels 1-3) to all transported mRNAs. The total SFARI group was also divided into the ones that are also in the FMRP CLIP group (17, 34) and the ones that are not. For all groups, there was a cut-off of 1RPKM to avoid mRNAs not being expressed at this point in development. P values from comparison to all mRNAs (Students t-test with Bonferroni correction for multiple tests (n=14 for all comparisons in figure). Only Significant P values (p<0.01 after correction are shown). The N for each comparison group is shown under the group.

**Supplemental Figure 6.**
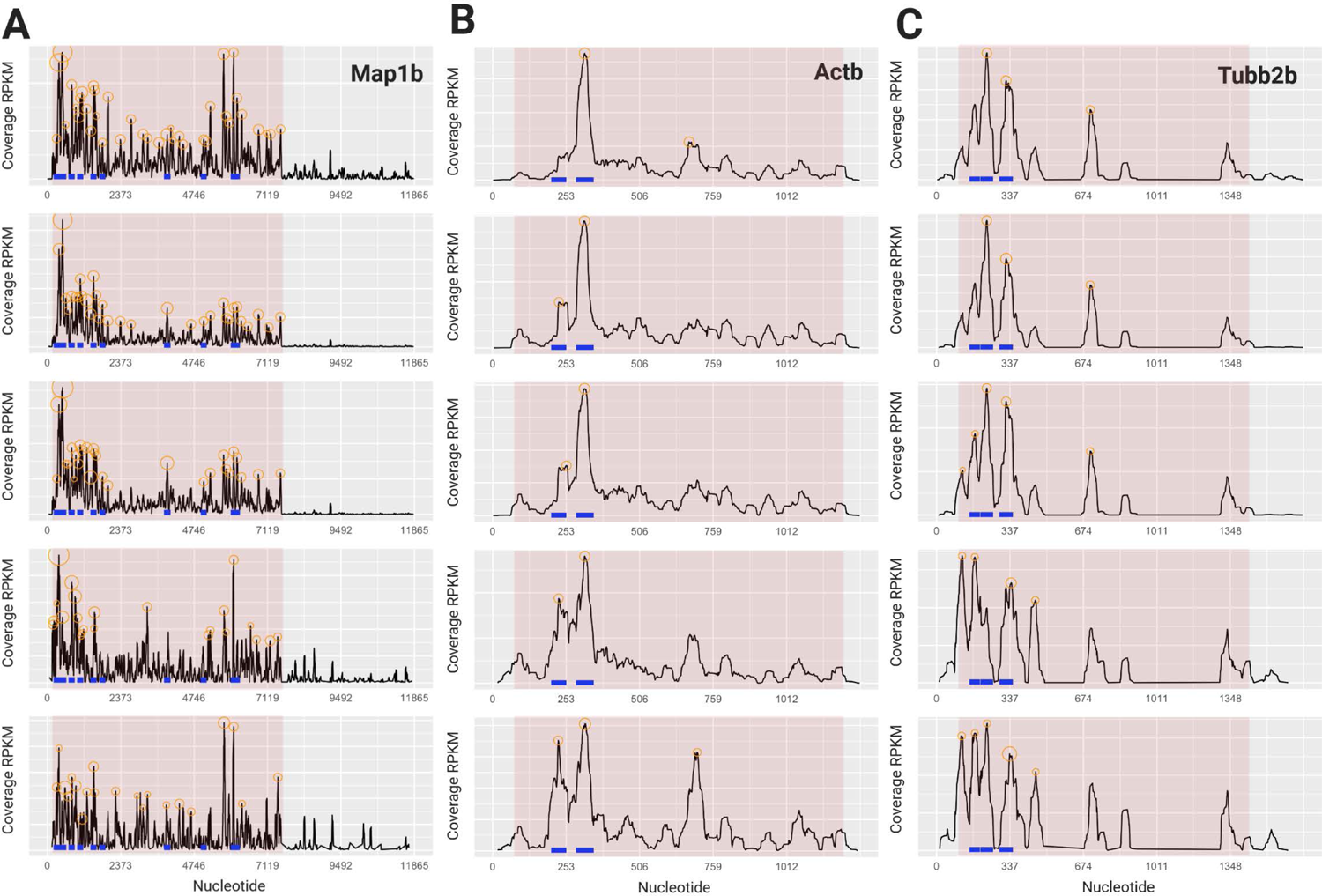
Read Maps of all 5 biological replicates. (A-C) Footprint read maps for Map1b (A), Beta-actin (B) and Tubulin 2b (C) for 5 biological replicates. Circles represent consensus peaks of footprint reads mapping to the same sequence, blue lines represent reproducible consensus sequences across biological replicates. Criteria for identifying consensus sequences (blue lines) is that at least 3 of 5 biological replicates have a consensus peak at this site. Red shading represents coding sequence (CDS).

**Supplemental Figure 7.**
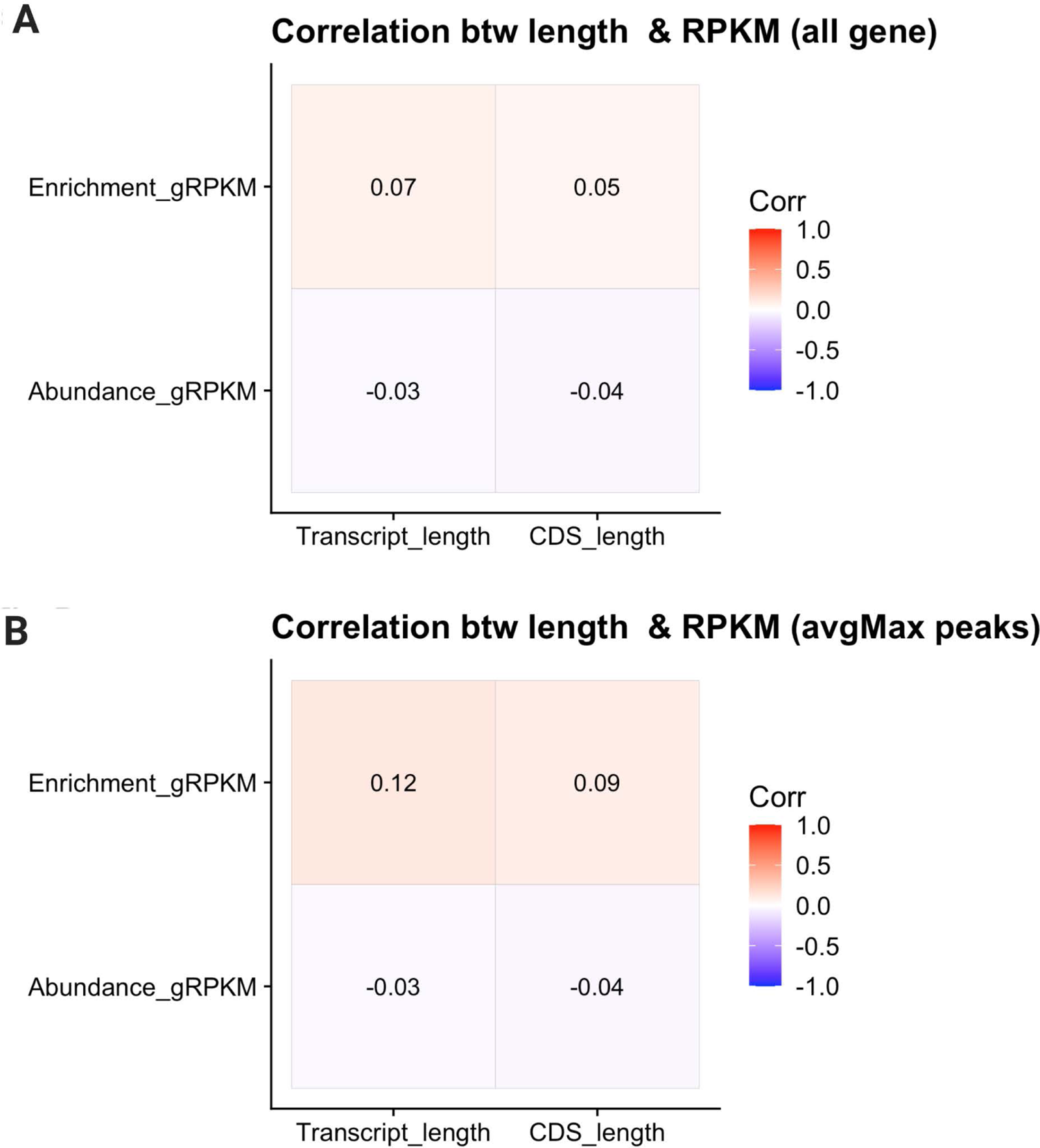
Lack of correlation between transcript length and transcript enrichment and abundance. The correlation with the length of the transcripts either the entire transcript length or the coding sequence (CDS) length with the amount of enrichment and abundance of the mRNA were calculated for either the total set of mRNAs (A) or just the mRNAs with peaks (B). R values are shown, and the squares colored based on the heat map of the R scores (inset). No correlation was significant (P<0.05).

**Data Set 1: Abundance and Enrichment for all mRNAs** Abundance and Enrichment of footprint reads. For each mRNA, the genename (colum A), the average RKPM from the RNA-SEQ of the total RNA Column E, n=5), the average RKPM of the footprint reads (Coliun F, n=5) are shown. The average RPKM was also independently calculated after first separating reads into long (>33, column G)), medium (25-33, column H) and short (<25, column I) reads. The enrichment was calculated as the log Fold Change (FC) between RKPM of the footprint reads and the RKPM of the total reads (column B). Calculaion by Llima of the p value for the fold change (column C) and the false discovery rate (Column D) are also shown. For subsets of mRNAs used in Figure 5, Columns in the Master sheet show which mRNAs are chosen for each group, and these groups and the data associated them shown in separate sheets. The master sheet has all mNRAs, but the separate sheets are restricted to mRNS with Footprint RKPMs of >5.

**Data Set 2. GO analysis of enriched transcripts.** Full GO analysis based on top enrichment transcripts (fold change of RPKM footprint abundance)s/RKPM total RNA abundance) as seen in Supplemental Table 1. The different sheets represent analysis based on the top 50, 100, 200 or 500 transcripts on the list. Analysis is from the Web site gProfileR (58) using all Rat genes as a comparison group. MF (molecular function); BP (biological process); CC (cellular component). The interactions (column J) represent the members of the enrichment list included in the GO term (column B).

**Data Set 3. GO analysis of abundant transcripts.** Full GO analysis based on most abundant transcripts (RPKM footprint abundance) as seen in Supplemental Table 1. The different sheets represent analysis based on the top 50, 100, 200 or 500 transcripts on the list. Analysis is from the Web site gProfileR (58) using all Rat genes as a comparison group. MF (molecular function); BP (biological process); CC (cellular component). The interactions (column J) represent the members of the abundance list included in the GO term (column B).

**Data Set 4. Peaks of mRNA footprint reads.** List of all Peaks from the large (>32nt) footprint reds. The transcript ensemble number is given in Column A; starting and ending nucleotide of the peak in Column B and C respectively. Length of the peak is in colun D. The start and end of the corresponding region (5’UTR, CDS or 3’UTR is given in columns E and F respectively. The genename is givein in Column D, the region of the peak, including peaks at the start and the stop are given in column H, Column I is the number of total WGGA and RGACH reads in each peak that contains both motifs. Column J gives the placement of the peak with respect to the normalized CDS. The number of sequences matching GVAGAW and GACAAG (based on FIMO) are given in columns K and L respectively. For sites with no shared WGGA and RGACH site, the number of WGGA sites and RGACH sites are given in columns M and N respectively. Column O is the sum total of the sites measured in I and K-0) for each peak.

**Data Set 5. RBP consensus sites in peaks.** The results of FIMO screening for RBP consensus sites (from ref 38) with additional FMRP sites from Ref 20 and 21). The RBP is given in colum A, the motif in column B and the number of peaks (from the large reads >32nt) with a match from the FIMO search (p<0.05) in column C. The total number of occurences of the motif (there may be muliple matchs in each peak) is given in D, and the distribution of the number of matches in each peak is given in E.

---

**Supplementary Table S1.**
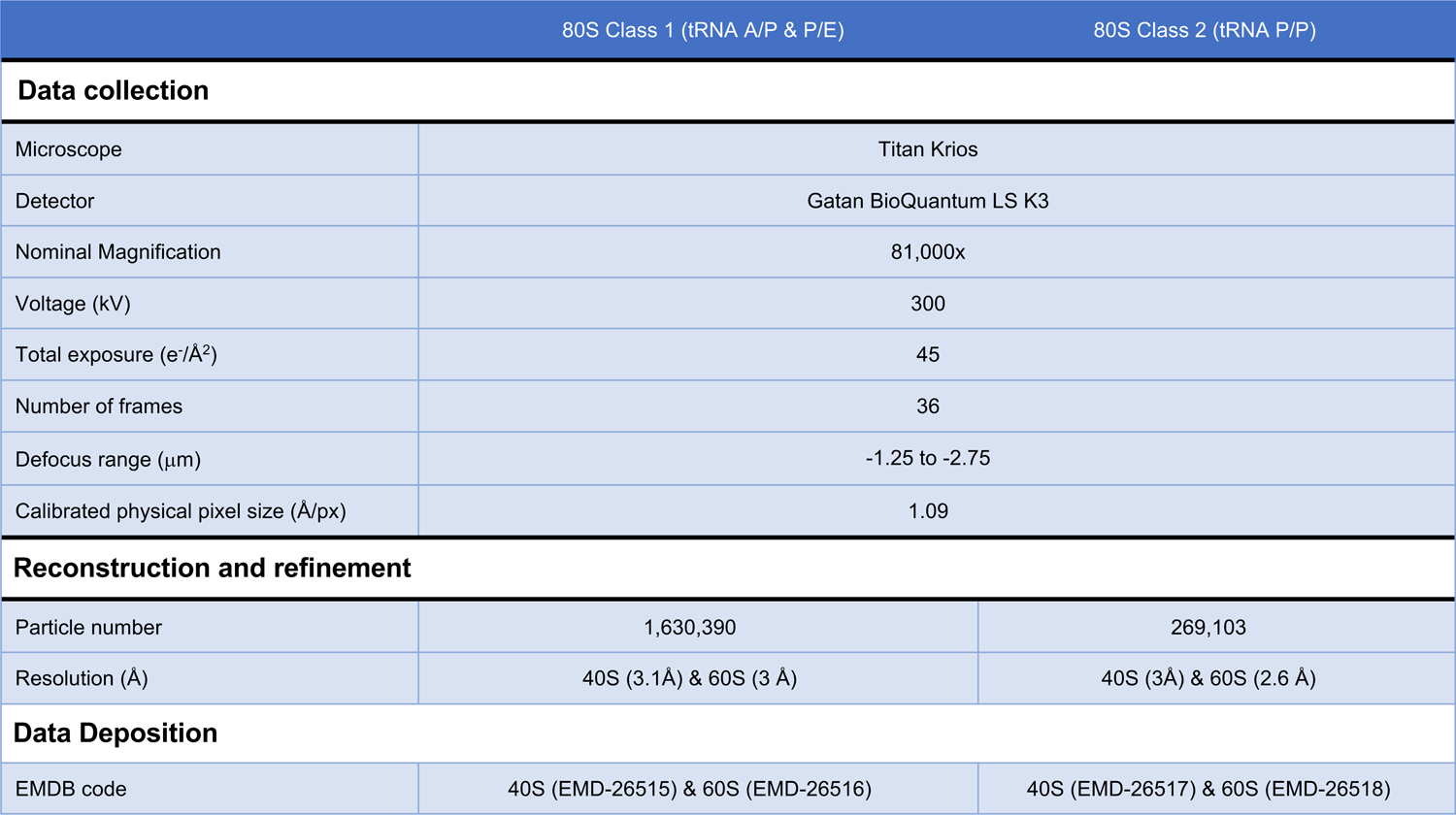
**Cryo-EM analysis of 80S ribosomes from RNA granules.** Data acquisition parameters, map statistics.

